# Molecular and Structural Characterization Reveals Divergent Extracellular Vesicle Profiles Between Wild Type and Alzheimer’s Disease Cerebrocortical Organoids

**DOI:** 10.64898/2026.05.13.724352

**Authors:** Anthony Balistreri, Natalie Turner, Jadon Compher, Mireya Almaraz, Akhil Prabhavalkar, Sachita Chittal, Sergio R. Labra, Kehinde Ezekiel, Christine Baal, Claudia Cedeño Kwong, Swagata Ghatak, Jan-Hannes Schäfer, Kimberly Vanderpool, Kathryn Spencer, John R. Yates, John P. Nolan, Scott Henderson, Stuart A. Lipton, Jeffery W. Kelly

**Affiliations:** Department of Chemistry, The Scripps Research Institute, La Jolla, CA 92037, USA; Department of Integrative Structural and Computational Biology, The Scripps Research Institute, La Jolla, CA 92037, USA; Department of Molecular & Cellular Biology, The Scripps Research Institute, La Jolla, CA 92037, USA; Core Microscopy Facility, The Scripps Research Institute, La Jolla, CA 92037, USA; The Scintillon Institute, San Diego, CA 92121, USA; Neurodegeneration New Medicines Center, The Scripps Research Institute, La Jolla, CA 92037, USA; Department of Neurosciences, School of Medicine, University of California at San Diego, La Jolla, CA 92093, USA

## Abstract

Alzheimer’s disease (AD) is a neurodegenerative disorder affecting millions of patients globally. Despite significant efforts from researchers in recent decades, there are still many unanswered questions about AD pathogenesis. AD patient brains manifest changes in extracellular vesicles (EVs) secreted from diseased neurons, and the effect of this phenomenon remains poorly understood. EVs contain a variety of biomolecules and play a critical role in cell-to-cell communication in all eukaryotic organisms. Here, we report a thorough characterization of small EVs purified from cultures of human cerebrocortical organoids. These organoids are differentiated from human patient-derived stem cells that bear a familial AD mutation in the presenilin 1 (PSEN1) gene, or from an isogenic wildtype (WT) control. The organoid conditioned media was aspirated from cultures and processed for EV enrichment using a non-invasive technique that requires no cellular disruption. EVs purified from AD organoid conditioned media have a wider size distribution and show differential expression of tetraspanins CD63, CD9, and CD81 when compared to WT organoid-derived EVs. AD organoid-derived EVs can have single, double, and even triple membranes and display luminal fibrillar material. A deep proteomic profiling of the EVs reveals several statistically significant differences, including evidence for modifications in secretory autophagy. EV isolates from both WT and AD organoids show strong binding to amyloid detecting dyes, both in bulk fluorescence and fluorescence microscopy assays. After a 1-week co-culture of AD organoids with WT organoids, there is evidence of endosomal membrane transfer between the isogenic cultures with an increase in amyloid-β peptides in the WT organoids. These observations support the notion that non-cell-autonomous spread of amyloid-containing EVs in human AD brains can be modeled in a cerebral organoid system.

## 1. INTRODUCTION

Alzheimer’s disease (AD) is a neurodegenerative disorder affecting 50 million patients worldwide, with predictions of increasing prevalence (Association 2025). The majority of AD cases are sporadic, where unknown aging-linked changes, which include environmental and/or genetic insults, burden protein homeostasis manifesting in cell death and cognitive impairment (Selkoe & Hardy 2016). A subset of patients develop inherited AD–their genomes contain a mutation(s) known to increase the risk of developing AD earlier than in sporadic cases (Selkoe & Hardy 2016). One example is the M146V mutation in PSEN1, giving rise to altered processing of amyloid precursor protein, causing neurons to produce higher amounts of the aggregation-prone amyloid-β (Aβ)_1_-_42_ peptide (Wang *et al*. 2006).

Recent advances in differentiation techniques have enabled modeling of neurodegeneration using complex 3D cultures (Faravelli *et al*. 2025). Our group recently published a human induced pluripotent stem cell (hiPSC)-derived 3D cerebrocortical organoid model of AD, comprising multiple cell types, including excitatory neurons, inhibitory neurons, and glial cells, displaying several measurable AD phenotypes (Labra *et al*. 2026). The M146V PSEN1 hereditary AD organoids differ from their isogenic, gene-corrected controls by exhibiting higher levels of Aβ peptides and phosphorylated tau while also displaying a clinically relevant AD neuronal hyperexcitability phenotype (Labra *et al*. 2026).

Reduced neuronal endolysosomal system (ELS) function, and modification of extracellular vesicles (EVs) secreted from affected cells are features of AD pathology. The ELS is a collection of endosomes that traffic cargo to and from various compartments in the cell; it is a critical part of maintaining cell homeostasis (Klumperman & Raposo 2014). The ELS is disturbed in many neurodegenerative diseases, including AD and Parkinson’s disease (Herman *et al*. 2024). Though the mechanism is not well understood, autophagy and lysosomal deficiency are implicated (Wolfe *et al*. 2013). Fortunately, we can study these phenomena by non-invasively gathering EVs from conditioned media, affording snapshots of the health of the tissues.

Small EVs, between 30 and 200 nm in diameter, play an important role in eukaryotic cell-to-cell communication (Raposo & Stoorvogel 2013). Exosomes, a class of small EVs, arise from multivesicular bodies (MVBs), dynamic intracellular membrane-bound vesicle organelles formed by the inward budding of the endosomal membrane during endosome maturation. MVBs are a key component of the ELS and function to traffic molecular cargo out of the cell or reroute ubiquitinated proteins to the lysosome (Raposo & Stoorvogel 2013). SNARE proteins and Rab GTPases mediate MVB fusion with the plasma membrane to release intraluminal vesicles as exosomes (Raposo & Stoorvogel 2013). The other common pathway of EV secretion is via budding of the plasma membrane (Jeppesen *et al*. 2023). Plasma membrane protrusions are pinched off via scission, and the resulting EVs are referred to as ectosomes or microvesicles (Jeppesen *et al*. 2023). Exosome and microvesicles are just two vesicle subtypes that are both distinct in their biogenesis and possess distinct molecular cargoes (Cocucci & Meldolesi 2015; Mathieu *et al*. 2021). However, the overlap in EV biophysical characteristics pose significant challenges to researchers seeking to define the unique properties associated with each subtype. Once secreted, EVs of all types can be readily endocytosed by neighboring cells, effectively delivering their cargo.

EVs can be isolated from many biofluids, and have many potentially effective uses in biomedical research, including for disease diagnosis (Colombo *et al*. 2014). Analysis of EV cargo is a valuable tool for understanding the intracellular protein dynamics of the cell-of-origin and can provide insight into disease mechanisms not readily distinguished from whole biofluid or tissue analysis (Kalluri & LeBleu 2020).

Here we developed a robust methodology to enrich EVs from cerebrocortical organoid conditioned media using a combination of size-based purification strategies. We subsequently used biochemical, biophysical, and analytical chemistry tools to comprehensively characterize the EVs secreted by hereditary AD and gene corrected organoids. We measured the count and size distribution of organoid-derived EVs using vesicle flow cytometry and performed structural biology studies and observed multilamellar vesicles in the AD organoid-derived EVs. Deep proteomic profiling of EV isolates revealed significant differences in protein cargo of AD vs. control organoid-derived EVs. We found that organoid derived EVs contain several amyloidogenic proteins and co-culturing WT organoids with AD organoids exacerbates amyloid pathology in the WT cells.

## 2. METHODS

### 2.1 hiPSC maintenance and cerebrocortical organoid differentiation

All hiPSC and cerebrocortical organoid tissue culture was performed in accordance with a previously established protocol (Labra *et al*. 2026). Briefly, isogenic hiPSC lines (WT and PSEN1^M146V^/WT familial AD mutant) derived from a male donor were obtained from the Tessier-Lavigne laboratory and New York Stem Cell Foundation (Paquet *et al*. 2016). Pluripotency and euploidy were confirmed by immunolabeling and G-banding, respectively. Cultures were maintained in parallel, regularly karyotyped and tested for mycoplasma, and passaged no more than 13 times.

For organoid generation, hiPSCs were dissociated with Accutase, aggregated in Aggrewell plates with mTeSR Plus + CEPT (Chroman I, Emricasan, Polyamine supplement, and Trans-ISRIB) for 24 h, then transferred to ultra-low attachment plates on an orbital shaker (60 RPM). Embryoid bodies were cultured in Essential 6/hESC medium with WNT/Dual SMAD inhibition for 6 days, then in Neural Media (NM) supplemented with EGF2/FGF2. From day 25–43, growth factors were replaced with neurotrophic factors BDNF and NT-3; organoids were then maintained long term in NM (Neurobasal™-A Medium, B-27™ Supplement, minus vitamin A, 1% GlutaMAX™ Supplement). Around day 35, organoids were moved to 100 mm ultra-low attachment dishes and kept shaking. Mature cultures were split as needed to maintain a confluency of approximately 15-20 organoids per plate and tested for mycoplasma every 1–2 months.

### 2.2 EV purification

EVs were purified using a two-step purification process. WT or AD organoids were cultured ultra-low attachment 10 cm dishes in NM. After 4 days, the medium was collected from 6 dishes containing approximately 40 individual organoids, totaling 120 mL of conditioned medium. The medium was concentrated using a TFF-Easy device (HBM-TFF/1) according to the manufacturer’s instructions. Briefly, the medium was passed through the center of the TFF-Easy device using a peristaltic pump set to at least 50 mL/min flow speed. The permeate sample was gathered through the permeate nozzles at the bottom of the device. The medium was circulated through the device until it concentrated down to 1 mL. The permeate and TFF concentrate were flash frozen in liquid N_2_ and stored at -80 °C until further use.

On the day of a given experiment, the TFF concentrate and permeate were thawed at room temperature. EVs were purified from the TFF concentrate using an Izon qEV 35nm column (ICO-35) according to the manufacturer instructions. Briefly, the qEV column was equilibrated with 5 CV of EV buffer (DPBS Ca2+ Mg2+, 14040133). The 1 mL TFF concentrate was added to the top of the column and 2 mL of EV buffer was added afterward. 2 additional mL of EV buffer was added and the eluate collected; this fraction contained the EVs. An additional 1 mL of EV buffer was added and the eluate collected; this fraction was the HMW protein sample. The 2 mL EV sample was concentrated using a 10K MWCO spin column (VS0102) and centrifuged at 4 °C at 10,000 x g for 15 min intervals until the desired volume was reached. All EV purification samples were used for analysis within 24 hours of qEV column purification.

### 2.3 Western Blot

SDS-PAGE and western blot analysis was performed according to previously established protocols (Labra *et al*. 2026). Briefly, the protein concentration of EV purification samples and organoid lysates was measured via Pierce BCA assay. The protein concentration was normalized to 12 µg of total protein for each lane except for the permeate lanes which were too low in protein concentration and remained undiluted. The samples were separated by SDS-PAGE and transferred to PVDF membranes by dry-transfer. After blocking in Intercept® Blocking Buffer, membranes were probed with the following primary antibodies: Calnexin (Rb, 1:1000, ab133615); FLOT1(Rb, 1:1000, JB19-45); CD9 (Rb, 1:500, ab263019), Abeta XP (Rb 1:1000, 8243S). Secondary antibodies used were IRDye® 680LT Goat anti-Rabbit (1:10,000 dil., 926-68021) and IRDye® 800CW Goat anti-Mouse (1:10,000 dil., 926-32210). Membranes were visualized using a LiCor Odyssey FC and ImageStudio software (v5.2).

### 2.4 Immunoprecipitation and ELISA

Protein G Dynabeads (10004D) were modified and immunoprecipitation was performed according to manufacturer’s instructions. Briefly, Protein G Dynabeads were prepared using amine-reactive crosslinking to immobilize a cocktail of 5 µg each of the following TS capture antibodies: CD63 antibody (ab134045), CD81 antibody (ab79559), CD9 antibody (ab263019). Final EV isolates and permeates were split in half and added to TS+ or unmodified (TS-) Dynabeads and incubated with tilting and rotation overnight at 4 °C. Using a magnetic tube rack, the beads were separated from the supernatant, which was collected and called the flow through. The beads were rinsed three times with DPBS Ca2+ Mg2+ until being treated with an EV lysis buffer (DPBS Ca2+ Mg2+, + 2% SDS) to release all EV associated proteins. The resulting eluate and flow through were diluted with DPBS Ca2+ Mg2+ or EV lysis buffer to produce 1% detergent concentration across all samples.

Total tau and Aβ_42_ levels in immunoprecipitations samples were quantified using the Invitrogen Human Total Tau ELISA Kit (Thermo Fisher Scientific, KHB0041) and Human Amyloid beta 42 ELISA Kit (Thermo Fisher Scientific, KHB3441) according to the manufacturer’s instructions. Briefly, samples were diluted 1:10 in standard diluent buffer and incubated in pre-coated plates. Absorbance was measured at 450 nm using a PHERAstar microplate reader. Concentrations were calculated using GraphPad Prism v10 after generating a standard curve using a 4-parameter algorithm.

### 2.5 Vesicle flow cytometry (vFC)

Vesicle concentration, size, and tetraspanin (TS)-positive fraction and were determined by single vesicle flow cytometry(Sandau *et al*. 2020; Tekkatte *et al*. 2023). using a commercial kit (vFC Assay kit, CBS-4; Cellarcus Biosciences, San Diego, CA) and flow cytometer (Cytek Aurora, TSRI Flow Cytometry core) that had been calibrated using a vesicle size (Lipo100, CBS-1) and fluorescence intensity standards (nanoCal, CBS-7). Briefly, samples were stained with the fluorogenic membrane stain vFRed and a mix of fluorescent antibodies targeting CD9, CD63, CD81 (vTag TS-APC mix, CBS-5), molecules often associated with some EVs, for 1h at RT, diluted and measured. Data were analyzed using FCS Express (De Novo Software) to gate membrane-positive events, estimate their size (surface area and diameter) and immunofluorescence (reported in units of antibody binding capacity, ABC). The assay design included a pre-stain dilution series to determine the optimal initial sample dilution and multiple positive and negative controls, per guidelines of the International Society for Extracellular Vesicles (ISEV) (Welsh *et al*. 2024).

### 2.6 Negative staining transmission electron microscopy (TEM)

Carbon-coated□copper grids (400 mesh, Ted Pella, Inc., 01844) were glow-discharged and□10□µL of each sample was adsorbed for two minutes. Excess sample was wicked away and grids were negatively stained with 2% uranyl= acetate (Ted Pella, Inc, 19481)□for 2 minutes. Excess stain was wicked away, and the grids were allowed to dry. Samples were analyzed at □120□kV with a□Thermo Fisher□Talos L120C transmission electron microscope and images were□acquired□with a CETA 16M CMOS camera.

### 2.7 Preparation of samples for cryo-EM

UltraAuFoil 200 mesh R 2/2 grids (Q250AR2A) were glow-discharged under vacuum for 30 s at 15 mA in a Pelco easiGlow 91000 glow discharge cleaning system (Ted Pella). 3.5 µl sample was applied to the front of the grid and incubated for 3 min at 4°C. After removal of excess sample with Whatman 1 filter paper (WHA1001090), 3.5 µl sample was applied to the front of the grid and 0.5 µl to the back of the grid. After 30 s incubation, grids were back-blotted with Whatman 1 filter paper for 4-6 s after the liquid spot on the filter paper stopped spreading. Grids were manually plunge-frozen in a 4 °C cold room with >95% humidity. AutoGrids were assembled and immediately screened.

### 2.8 Cryo-EM data collection and processing

Cryo-EM datasets were collected on a Talos Arctica TEM (Thermo Fisher) operating at 200 keV. Movies were recorded using a Falcon 4i direct electron detector (Thermo Fisher) at a nominal magnification of 120,000, corresponding to a pixel size of 1.2 Å. Movies were saved in the electron-event representation (EER) format and recorded at a total electron exposure of 50 e^−^ per Å^2^. All datasets were collected automatically using EPU (v.3.9, Thermo Fisher) with a defocus range of −1.0 to −3.0 μm. EPUs fast exposure navigation was used to collect data up to an image shift of 8 µm. All datasets were processed using cryoSPARC (v.4.7). Movies were collected and dose-fractioned into 40 frames, motion corrected using patch-based motion correction, followed by patch-based CTF estimation in cryo-SPARC live.

### 2.9 Cryo-TEM analysis of EVs

Micrographs were manually inspected for EVs. The wildtype-dataset showed vesicles on 17 of 179 micrographs and the AD-dataset was positive for vesicles on 63 out of 431 micrographs. Images were low-pass filtered to 2 Å and binned by four to 1024×1024 px .png images before vesicle-area determination in *Fiji* (Schindelin *et al*. 2012). Violin-plots of EV areas were generated with custom python scripts using *matplotlib*. Pairwise significance was computed with Mann-Whitney U test using *scipy*.

### 2.10 Proteomics

EV-enriched samples were prepared for proteomic analysis using a modified filter aided sample preparation protocol for EV proteomics, with all buffers, reagents, and centrifugation steps performed as previously described (Turner *et al*. 2022; Wiśniewski *et al*. 2009). In brief, EV-enriched samples were subjected to acetone protein precipitation and resuspended in a small volume of 1% w/v sodium dodecyl sulfate. Protein extracts were mixed with a lysis buffer containing 1% w/v sodium deoxycholate and protease inhibitors (Pierce Protease Inhibitor Cocktail Mini Tablets, Thermo Scientific) to a final concentration of 0.5% (1:1 sample to buffer ratio). Samples were sonicated in a water bath for 5 min and incubated on ice for 15 min, before being loaded onto 30 K centrifugal filter columns (Nanosep Omega, Cytiva, cat no OD030C35). Samples were centrifuged at 14,000 xg for 15 min at 21 °C, or until the sample volume had completely passed through the filter. Proteins bound to the filter were reduced with dithiothreitol (DTT) for 1 h at RT and columns centrifuged again, and a column wash performed with 8 M urea. Alkylation of cysteine (C) residues was performed by incubating samples with iodoacetamide for 20 min at RT in the dark. Columns were centrifuged again and washed twice with 8 M Urea and twice with 100 mM ammonium bicarbonate (AMBIC). Trypsin suspended in 100 mM AMBIC was added to the filters at an enzyme:protein ratio of 1:50, and columns were incubated in a humidified chamber at 37 °C overnight. The next day, peptides were recovered by centrifugation into a clean 1.5 mL protein lobind microcentrifuge tube (Eppendorf), and one additional elution performed with 40 µL 100 mM AMBIC. Peptide concentrations were quantified by colorimetric peptide assay (Pierce, Thermo Scientific), and equal amounts of peptides loaded onto Evotips (Evosep) in triplicate, according to the manufacturer’s instructions.

### 2.11 Liquid chromatography tandem mass spectrometry (LC MS/MS)

MS data were acquired on a timsTOF Pro2 mass spectrometer (Bruker) equipped with electrospray ionization source, coupled to an Evosep nano-LC system. The instrument was operated in data-independent acquisition with parallel accumulation serial fragmentation (DIA PASEF) mode (Meier *et al*. 2015; Venable *et al*. 2004) and samples were run with a 15 spd (∼88 min) method with a flowrate of 220 nL/min. Buffer A consisted of H_2_O/0.1% FA and Buffer B was Acetonitrile/0.1% FA (LC-MS grade, Fisher Scientific). Peptides were separated by reversed-phase HPLC on an in-house packed analytical capillary column with integrated in-house pulled tip, with dimensions 25 cm, 150 nm internal diameter, and BEH 1.7 µm C18 resin (Waters). Capillary voltage was set to 1500 V and dry gas flow and temperature 3.0 L/min and 180 °C, respectively. DIA-PASEF MS1 scan range was 100 – 1700 *m*/*z* and ion mobility (1/K0) = 0.6 – 1.6 V s cm^2^. Accumulation and ramp time were 100 ms, and cycle time was 1.16 s. The PASEF MS/MS window scheme was set to 32 wide *m/z* isolation windows from 400 – 1200 *m/*z of width 25 *m*/*z* with 1 *m*/*z* overlap. MS2 TOF resolution was 45,000 and singly charged precursor exclusion enabled. Five-hundred ng of peptides per sample were injected into the mass spectrometer in technical triplicate via elution from the column tip into a captive spray source, and peptides were nanosprayed into the MS.

### 2.12 Proteomics data analysis

MS data files (d folders) were loaded in DIA-NN v2.2.0 and searched against a predicted *Homo sapiens* library containing 20,405 reviewed sequences (downloaded from Uniprot: 22 Oct 2024) at 1% FDR. Trypsin was set as the digestion enzyme, carbamidomethylation on C was set as a fixed modification, with up to 3 variable modifications and 1 missed cleavage allowed. All other settings were as default and MBR was disabled. The output report file was imported into RStudio for further processing using the ‘MSstats’ ( v4.16.1) and ‘MSstatsConvert’ packages (v1.19.3 development version, available from Bioconductor: https://www.bioconductor.org/packages/devel/bioc/html/MSstatsConvert.html) (Kohler *et al*. 2023). Data were filtered for common contaminants included in the .fasta file from the Cambridge Centre for Proteomics (CCP) cRAP database (https://github.com/CambridgeCentreForProteomics), with additional filtering at *q* value, protein group *q* value and global *q* value ≤ 0.01. Settings in MSstats dataProcess were as follows: logTrans = 2, normalization = “equalizeMedians”, featureSubset = “all”, remove_uninformative_feature_outlier = TRUE, min_feature_count = 2, summaryMethod = “TMP”, censoredInt = “NA”, MBimpute = FALSE, remove50missing = TRUE. All other settings were as default. Aggregated protein quantities used for the PCA plot (mixOmics; v 6.32.0) (Rohart *et al*. 2017) and Venn diagram (VennDiagram; v1.7.3) were generated using the R package diann (v1.0.1). For the Venn diagram, data were filtered to include proteins identified in ≥2 out of 3 replicates within groups. The EnrichR (Kuleshov *et al*. 2016)package was used to perform enrichment analysis of proteins against the following databases: GO_Biological_Process_2023, GO_Molecular_Function_2023, GO_Cellular_Component_2023.

### 2.13 Thioflavin T (ThT) binding assay

The protein concentration in EV purification samples was assessed by Pierce BCA assay (23225). EV purification samples were diluted to 20 ug of protein in the wells of a 96 well plate (P96-1.5P) with DPBS Ca2+ Mg2+ and 10 µM Thioflavin-T (ThT, EW-88226-62, frozen stock stored at 1 mM in water). Any sample under 20 µg of protein was undiluted. Lipo100 was diluted to 25 x 10^5^ total vesicles per well. ThT fluorescence was monitored in a CLARIOstar Plus microplate reader by measuring fluorescence (excitation: 440nm, emission: 485nm) for 24 hours at 37 °C every 5 mins with a 5 sec shake before each reading. The assay was performed with three technical replicates and a ThT only baseline condition was subtracted from each average.

### 2.14 Stimulated emission depletion (STED) microscopy

EV purification samples were mixed with 10 ug/mL of AmyTracker 680 (A680-A-100) diluted in DPBS Ca2+ Mg2+ and spotted onto poly-L-lysine coated glass coverslips (CLS354085) and incubated at room temperature in the dark for at least 30 mins. The glass coverslips were mounted onto microscope slides and image data was acquired using an Abberior STED Facility Line microscope with an Olympus UPLXAPO 60x, NA 1.42 objective. Using 21.7% 561nm excitation laser, single plane STED image parameters were 30 nm^2^ pixels, 5 µsec pixel dwell time, 20 accumulations, with a 775nm depletion laser at 10%, and an emission band of 632 nm – 742 nm. Puncta size approximation was performed by opening the image in Fiji and drawing a straight line across a representative punctum and the Plot Profile tool was used to extract pixel intensity values. The resulting profile was fitted to a Gaussian function using Fiji’s (v154p) default curve-fitting routine. The full width at half maximum (FWHM) was calculated from the fitted parameter σ using: FWHM=2.35 × σ

### 2.15 Organoid co-culture and live-cell imaging

AD organoids were incubated overnight in NM supplemented with 2 µM Aco-650 (Acoerela). After incubation, the stained and an additional unstained organoid from the same batch were incubated for 10 mins in NM supplemented with the Hoechst nuclear stain (1:4000 dil., 62249). After several rinses, the organoids were moved to BrainPhys™ Imaging Optimized Neuronal Medium (05796) and placed in an optical bottom 96 well plate (07000629). The organoids were imaged using a Zeiss Celldiscoverer 7 microscope controlled by ZEN Blue (v3.13) with the specimen chamber set to 37 °C and CO_2_ 5%. Images were acquired with a 20×/0.7 NA objective in z-stacks spanning 10 µm. For each region of interest, a 2×2 grid (4 tiled fields) was captured using LSM Plus spectral multiplexing. Post-acquisition processing was performed in ZEN Blue using the LSM Plus module, Extended Depth of Focus method to make maximum image projection, and Stitching method using the Hoechst channel as a reference.

The Aco-650 stained AD organoids were placed in the same well of a 6-well plate with a WT organoid and co-cultured for 7 days, with fresh media swapped out every other day. To identify the WT organoid after co-culture, all WT organoids were prestained with CellTracker Green (1µM diluted in NM, C2925) for 30 mins. On the 7^th^ day, all organoids were removed from co-culture, briefly stained with Hoechst nuclear stain, and live-cell imaged using the above method.

### 2.16 Tissue clearing and immunohistochemistry

After live-cell imaging, organoids were fixed with 4% Paraformaldehyde (diluted from 16% PFA, AA433689M) at 4 °C overnight. Fixed organoids were rinsed with 0.3 M Glycine to quench any residual PFA. After two additional PBS rinses, the organoids were cleared following standard tissue clearing steps according to the SCALEVIEW-S4 (194-18561) protocol. Briefly, the fixed organoids were placed in SCALEVIEW-S4 for up to 72 hours at 42 °C with shaking at 800 rpm to remove occluding lipid. Cleared organoids were moved into blocking buffer (0.3% TBST + 3% NGS, 50197Z) and incubated for 1 hr at room temperature on a rocking shaker. Cleared organoids were incubated with the primary antibody solution (Blocking buffer + primary antibody) for 24 hours at 4 °C on a rocking shaker. The primary antibodies used were Abeta XP (1:500 dilution, Rb), AT8 (1:500 dilution, Mu), and MAP2 (1:1000 dilution, Ch). After rinsing with PBS, the organoids were incubated with secondary antibody solution (Blocking buffer + secondary antibody) for 4-6 hours at room temperature on a rocking shaker. The secondary antibodies used were Alexafluor+ 555 (1:2000, Goat anti Rb), Alexafluor+ 647 (1:2000, Donkey anti Ms), Alexafluor+ 488 (1:2000, Goat anti Ch). After washing out the secondary antibodies with PBS, the immunostained organoids underwent a second PFA fixation to fix the antibodies in place. After additional glycine and PBS rinses, the organoids were moved into an optical bottom 96 well plate and incubated in a mixture of 1:1 EasyIndex: DI water and Hoechst nuclear stain (1:4000 dil.) and incubated at 37 °C with shaking for 1-3 hours. Afterwards the solution was aspirated and undiluted EasyIndex (EI-100) was added to the organoids. Organoids reached transparency after a few additional hours and remained in Easy Index for imaging.

Cleared and immunostained organoids were imaged using a Zeiss Celldiscoverer 7 microscope controlled by ZEN Blue. Images were acquired with a 20×/0.7 NA objective in z-stacks spanning 20□µm. For each region of interest, a 3×3 grid (nine tiled fields) was captured using LSM Plus spectral multiplexing. Post-acquisition processing was performed in ZEN Blue using the LSM Plus module.

### 2.17 Image analysis

Processed image stacks were exported to Arivis Pro (v4.20) for quantitative analysis. Nuclei were segmented using the Cellpose Segmenter with the nucleus model, and Aβ signal was detected using the Simple Threshold analysis operator. Outliers were removed (3 outliers) using the ROUT method (Q = 1%). Additional image cropping and modification (e.g. adding scale bar, uniformly adjusting brightness and contrast settings across channels, etc.) for clarity of illustration was performed in Fiji v154p.

### 2.18 Quantification and statistical analysis

Graphs were produced in GraphPad Prism v10 and data shown is the average of *n* number of technical replicates (as noted in figure legend) with error bars depicting the standard error of the mean. Statistical significance was assigned based on a two-way ANOVA analysis using Prism’s default parameters where * denotes *p* < 0.05, ** denotes *p* < 0.005, *** denotes *p* < 0.0005, and **** denotes *p* < 0.00005.

## 3. RESULTS

### 3.1 Cerebral organoid conditioned media contains EVs

Human organoids are 3D cultures differentiated from hiPSCs, comprising multiple cell types that show promise for modeling human disease (Faravelli *et al*. 2025). In a recent publication, our lab produced a protocol for generating human cerebrocortical organoids (Labra *et al*. 2026). Specifically, we generated organoid pairs, one hiPSCs–organoid pair harboring a heterozygous familial AD mutation, e.g. PSEN1 M146V/WT used herein, along with an isogenic hiPSCs–organoid pair that have been gene corrected to WT to serve as a control. As a standard part of organoid differentiation and maintenance, the organoids are cultured over the course of months in serum-free growth medium. When the cultures are fed, the conditioned media is normally aspirated and discarded into liquid waste. Alternatively, we found that the conditioned media can be used as a plentiful source of EVs, which after purification can provide important molecular information about the organoids they came from through their characterization.

Eight-to-ten-week-old WT and AD organoids were incubated in culture medium for 4 days, after which 120 mL of conditioned media was pooled and concentrated 120-fold using tangential flow filtration (**Figure 1**). The tangential flow filter (TFF) yielded two solutions: i) a concentrate that was retained in the filter and contained EVs, and ii) an EV-depleted medium called a “permeate” that passed through the filter (**Figure 1**). The TFF concentrate was loaded into a size exclusion column to separate EVs from other high molecular weight contaminants (**Figure 1**). The size exclusion column afforded the final purified EV isolate. As an added control, we collected the subsequent elution fraction from the size exclusion column that contained high molecular weight (HMW) proteins and trace EVs. Purified EV isolates, permeates, and/or HMW protein samples were then subjected to a multitude of biochemical and analytical analyses (**Figure 1**).

**Figure 1.**
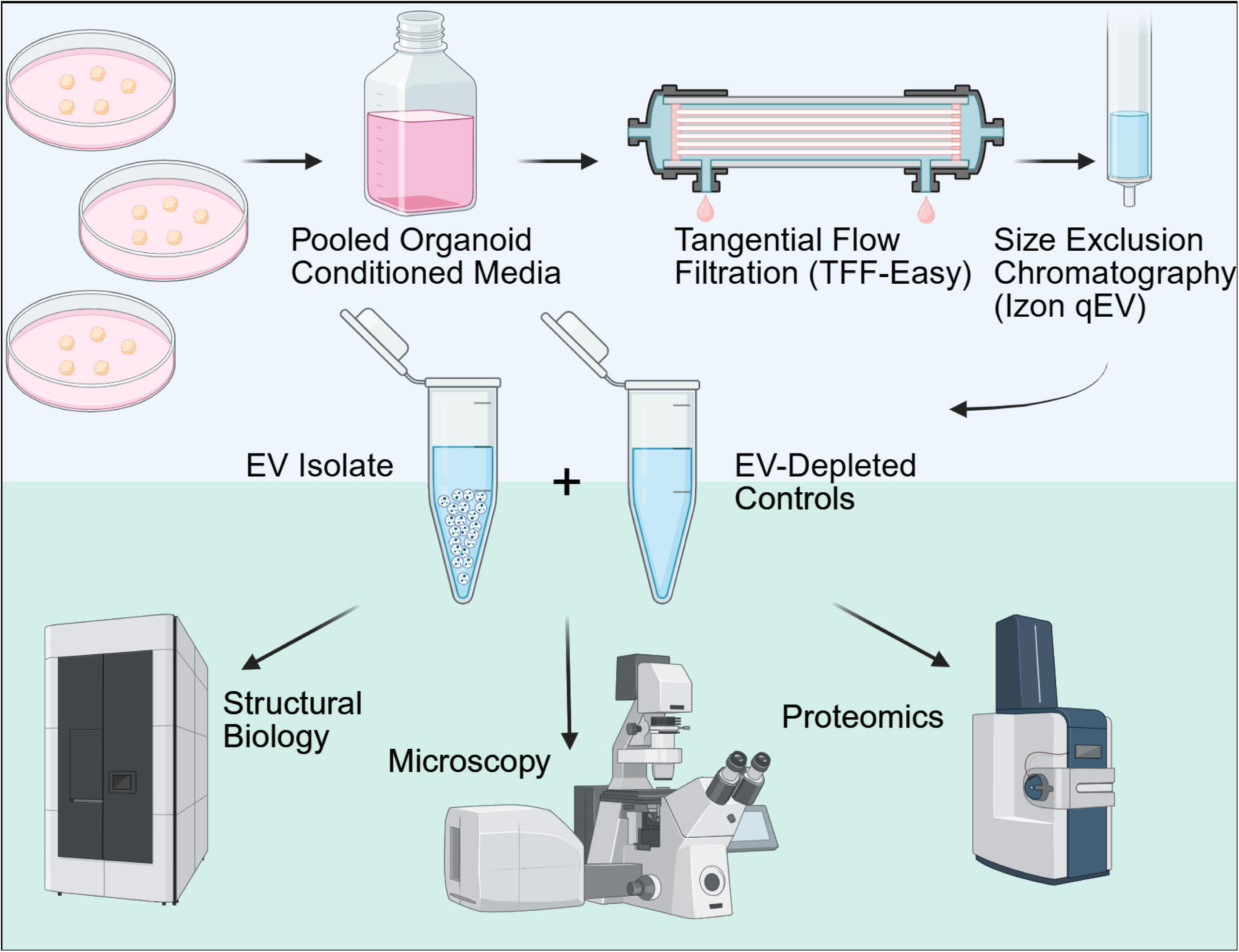
Purification method and applications workflow. EVs were purified through a two-step process that started with pooling organoid conditioned media from the same differentiation batch. Pooled conditioned media was concentrated 120-fold using tangential flow filtration. The TFF concentrate was separated by size exclusion chromatography and the final EV containing fraction. Three samples gathered during purification were analyzed further by biochemical and analytical methods.

### 3.2 EV isolates contain vesicles with standard morphologies and detectable EV surface marker proteins

WT and AD organoid conditioned media were processed as described above and the final EV isolate, the HMW protein sample, and the EV-depleted permeate were analyzed by Western Blot to detect EV markers (**Figure 2A**). A tetraspanin protein CD9 and Flotillin 1 are both markers of extracellular vesicle membranes (Kowal *et al*. 2016). CD9 and Flotillin 1 were both enriched in the EV isolates, compared to other fractions (**Figure 2A**). Calnexin, a common negative control for EVs expressed in the endoplasmic reticulum, was highly visible in the organoid lysate, but not in the EV isolates (**Figure 2A**). Interestingly, the Aβ XP antibody, which detects a variety of Aβ peptides by binding an epitope at the peptides’ amino terminus, produced a common band at approximately 40 kDa which could represent a stable Aβ oligomer or a structural component thereof that is SDS-resistant (**Figure 2A**)(Li *et al*. 2025). The full Aβ XP blot showed low molecular weight bands consistent with monomeric Aβ in the lysates and EV isolates (**Supplementary Figure 1**). Given the inconsistency of immunoblots in the detection Aβ from biological samples (Adlard *et al*. 2014), finding Aβ in the EV isolates was an initial indication that EVs could be an interesting analyte to study the abberrant Aβ processing that occurs in AD.

**Figure 2.**
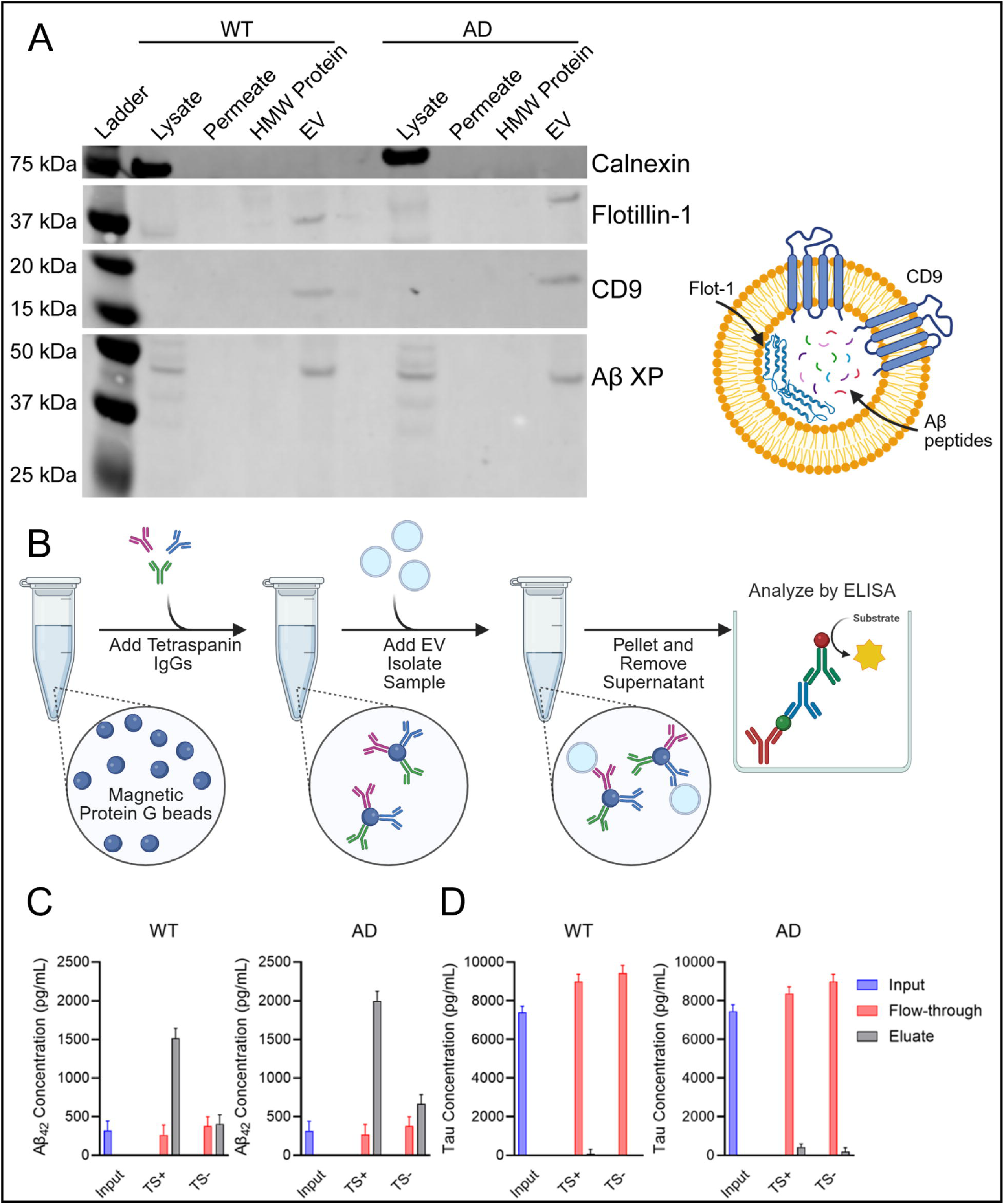
EV purification samples isolated from cerebrocortical organoid conditioned media contain EV markers, amyloid-β, and tau. **A)** A Western blot was performed on organoid lysate and EV purification samples to detect the presence of common EV markers and amyloid-β. Calnexin is an ER marker protein and was used as a negative control. Flotilin-1 and CD9 are two common proteins used as markers of EV membranes. **B)** A schematic showing the method of immunoprecipitation to further purify TS+ vesicles using a cocktail of TS antibodies (CD9, CD63, and CD81) from an EV-depleted media control (Permeate, **Supplementary Figure 2**) and final EV isolates. Sandwich ELISAs were used to detect **C)** Aβ42 and **D)** tau (total) in IP samples from EV isolates. After IP, proteins that did not bind to the beads were labeled Flow-through and proteins associated with the beads that were eluted from the beads using a detergent wash (1% SDS) were labeled Eluate.

Two significant AD proteins are enriched in both WT and AD organoid-derived EV isolates. Dysregulation and modified processing of amyloid precursor protein and tau are hallmarks of AD. Amyloid-β peptides were first observed to be associated with EVs in 2D HeLa and N2a cell culture media by Rajendran et al. (Rajendran *et al*. 2006) and tau was observed by Saman et al. in M1C cells and in the CSF of AD patients (Saman *et al*. 2012). After purifying EVs from our 3D culture media, an additional enrichment step was performed to confirm the presence of Aβ_42_ and tau in our EV isolates. Magnetic Protein G beads were modified with a trio of capture antibodies that bind to three common tetraspanin (TS) EV surface proteins: CD63, CD81, and CD9 (**Figure 2B**). The EV isolates and EV-depleted permeates from the WT and AD organoids were incubated with TS+ or TS- beads (a condition where the beads were not modified with capture antibodies). After incubation, the material that did not bind to the beads, the flow through, was separated from the beads. After several washes, the beads were treated with an EV lysis buffer and the resulting solution, the eluate, was collected (**Figure 2B**). Sandwich ELISAs were performed on all samples to detect two AD related proteins: Aβ_42_ (**Figure 2C, Supplementary Figure 2A**) and tau (**Figure 2D, Supplementary Figure 2B**). Aβ_42_ was present at the highest level in the eluate in the TS+ fractions, suggesting an enrichment associated with intact EVs with TS surface proteins **(Figure 2C**). Tau was present in predominantly the flow through, suggesting that tau is either co-purifying with the EVs or tau is associated with EVs that do not have TS proteins on their surface (**Figure 2D**).

Next, we used a calibrated flow cytometer and validated assay to measure EV concentration, size, and surface marker expression. The vFC assay employs a fluorogenic membrane probe, vFRed, to selectively stain EVs in proportion to their surface area, and validated antibodies to measure EV surface antigens. The Cytek Aurora flow cytometer was calibrated with standard beads to report immunofluorescence in standardized units of antibody binding capacity (ABC) and a synthetic vesicle standard to report vesicle size in units of surface area and equivalent diameter (assuming a spherical shape). Assay controls included buffer-only, reagent-only, and detergent treatment, as well as dilutions series to determine assay dynamic range and absence of particle coincidence (“swarm”) as described in the recent MISEV(Welsh *et al*. 2024) and MIFlowCyt-EV (Welsh *et al*. 2023) guidance documents (**Supplementary Figure 3**).

We found that EV isolates from WT (**Figure 3A-C**) and AD organoids (**Figure 3D-E**) contained vesicles ranging from the size limit of detection (LOD, ∼85 nm) to ∼ 270 nm (95^th^ percentile) with a median diameter of ∼124 nm. The AD organoid EV isolate contained more EVs compared to the WT organoid (∼3.1 e^9^ EVs/mL in the concentrated media compared to 1.6 e^9^/mL for the WT) but the EVs were phenotypically similar, exhibiting very low levels of tetraspanin expression (>20 ABC per EV; **Figure 3 B-C, E-F**) compared to TS-positive control EVs purified from human platelets (∼80 ABC per EV; **Supplementary Figure 3**). Tetraspanin positivity can be an indicator of vesicle biogenesis (Andreu & Yáñez-Mó 2014) and low positivity would indicate that EVs arising from MVBs make up a smaller proportion of the vesicles in the EV isolates compared to other classes of EVs.

**Figure 3.**
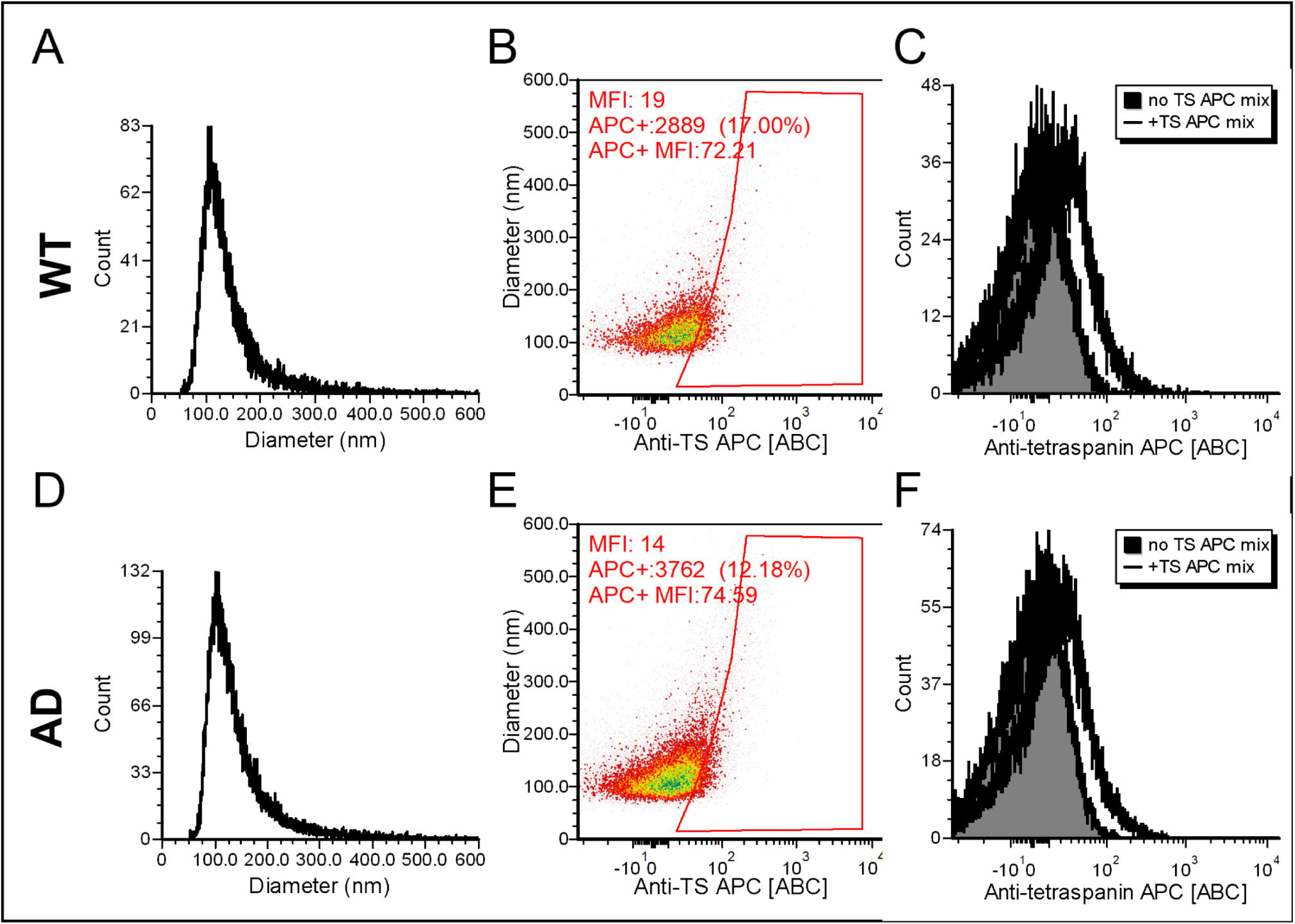
EVs isolates from AD and WT organoid media contain vesicles with low levels of tetraspanin expression. EV isolates were stained with a fluorogenic membrane probe and a trio of tetraspanin (CD9, CD63, CD81) binding antibodies conjugated to the APC fluorophore. Stained vesicles were flowed past a detector which measured the size and mean fluorescence intensity (MFI) of APC positive vesicles. EVs in media from WT (top) and AD (bottom) organoids were similar in size (∼85-270 nm; median 124 nm, **A and D)** and exhibited low levels of anti-TS antibody binding above background fluorescence **(B-C, E-F).**

### 3.3 TEM of EVs

Transmission electron microscopy is a common tool for visualizing vesicles in an EV isolate (Théry *et al*. 2006). Each of our EV purification samples was visualized by TEM. Cup-shaped vesicles of the expected diameter (∼30-200 nm) were identified in the EV samples, but not the HMW protein or permeate samples (**Figure 4A-F**). Though TEM is a powerful tool, it only provides a snapshot of the vesicles that adsorbed onto the TEM grid. Additionally, desiccation on the TEM grid leads to morphologies that give EVs a sunken, cup-like shape (**Figure 4B and C**) (Royo *et al*. 2020). Vitrification in solution by dipping TEM grids in liquid ethane solves the issue of non-physiological morphologies when visualizing EVs by preserving their natural shape in solution (Arraud *et al*. 2014). Cryo-TEM of our samples produced images of EVs that looked spherical (**Figure 4G and I**). Indeed, WT organoid-derived EVs were mostly spherical and homogeneous, though they did contain vesicles with different luminal densities (**Figure 4G**). On the other hand, AD organoid-derived EVs showed a diverse morphology, and they trended towards a wider, though not significantly different, size distribution (**Figure 4H and I**). There was a variety of multilamellar vesicles (MLVs) found in the AD organoid-derived EVs that were not as common in the WT organoid-derived EVs. Some vesicles had single, double, or triple membranes, suggesting a heterogeneous population of endosomes as the progenitors of the EVs (**Figure 4I**). Strikingly, large vesicles found in the AD organoid-derived EVs contained luminal fiber-like structures (**Figure 4I**). Additional fibers could be found nearby vesicles or in dense clumps that co-purified with the AD EVs (**Figure 4I**). The free-floating fibrils may result from co-purification and transient binding to EVs or be released after EV isolation owing to vesicle rupture (amyloid fibrils are established to mediate the rupture of membranes) (Jakhria *et al*. 2014). No such fibrils were observed in the WT EVs.

**Figure 4.**
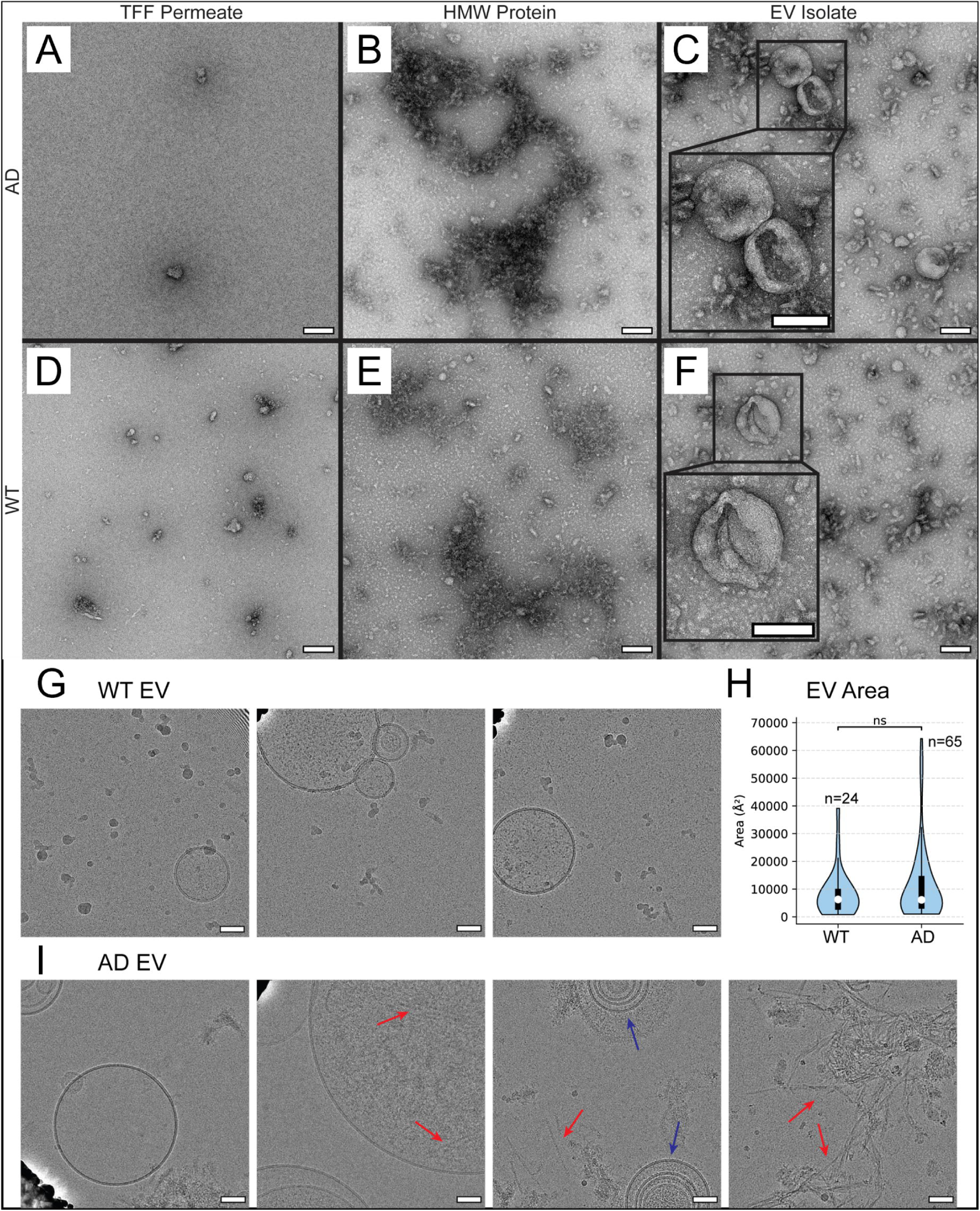
EVs isolates from AD and WT organoids contain vesicle-shaped structures that have divergent morphologies. Organoid conditioned media was processed to isolate EVs yielding **A and D)** an EV-depleted Permeate, **B and E)** a High MW Protein sample, and **C and F)** a final EV isolate. The samples were adsorbed onto carbon formvar grids, negatively stained with uranyl acetate, and visualized by TEM. The scale bars in all images represent 100 nm. Morphological analysis of WT and AD organoid derived EVs was also carried out with cryo-TEM**. G)** Micrographs of wildtype EVs. **H**) Vesicular area distribution for 24 WT EVs and 65 AD EVs. **I)** Micrographs of AD EVs. Scale bar 50 nm, multilamellar vesicles indicated with blue arrows, fibrils or amyloids indicated with red arrows.

### 3.4 Quantitative and qualitative proteomics analysis

Since EV protein content can change with the disease state of the secreting cells (Colombo *et al*. 2014), we sought to identify any significant differences between the proteins that were present in our WT and AD EV isolates. A total of 3848 proteins were quantified in WT and AD EVs samples (**Figure 5A**). Of these, 170 proteins were significantly upregulated in WT EV, and 184 proteins were upregulated in AD EV (FC ≥ 2, adj *p* ≤ 0.05). As it is currently unknown how AD pathology may affect the total populations of different vesicle classes (i.e. exosomes, microvesicles, etc.), we assessed the quantity of established EV protein markers (Welsh *et al*. 2024) between WT and AD organoid EV samples. CD9, SDCB1, FLOT2, and to a lesser extent TSG101 and FLOT1, were upregulated in AD EVs. Conversely, the validated neuronal EV marker L1CAM (Nogueras-Ortiz *et al*. 2024) was increased in WT EVs compared to AD EVs. L1CAM generally correlates with neuronal cell number, especially in early nervous system development (Barman & Thakur 2024; Patzke *et al*. 2016). The relative reduction observed may be indicative of early neuronal cell death or growth arrest in the AD organoids, which agrees with previous organoid characterization (Labra *et al*. 2026). These results suggest that AD pathology influences the proportions of vesicle subpopulations, and that the predominant cell-type-of-origin of vesicle populations may differ in disease states. This observation aligns with the hypothesis that AD affects the endolyosomal pathway, which is central to both lysosomal degradation and exosome biogenesis through MVB formation. The finding that EV subpopulations may be influenced by AD implicates secretory pathway involvement in disease pathogenesis.

**Figure 5.**
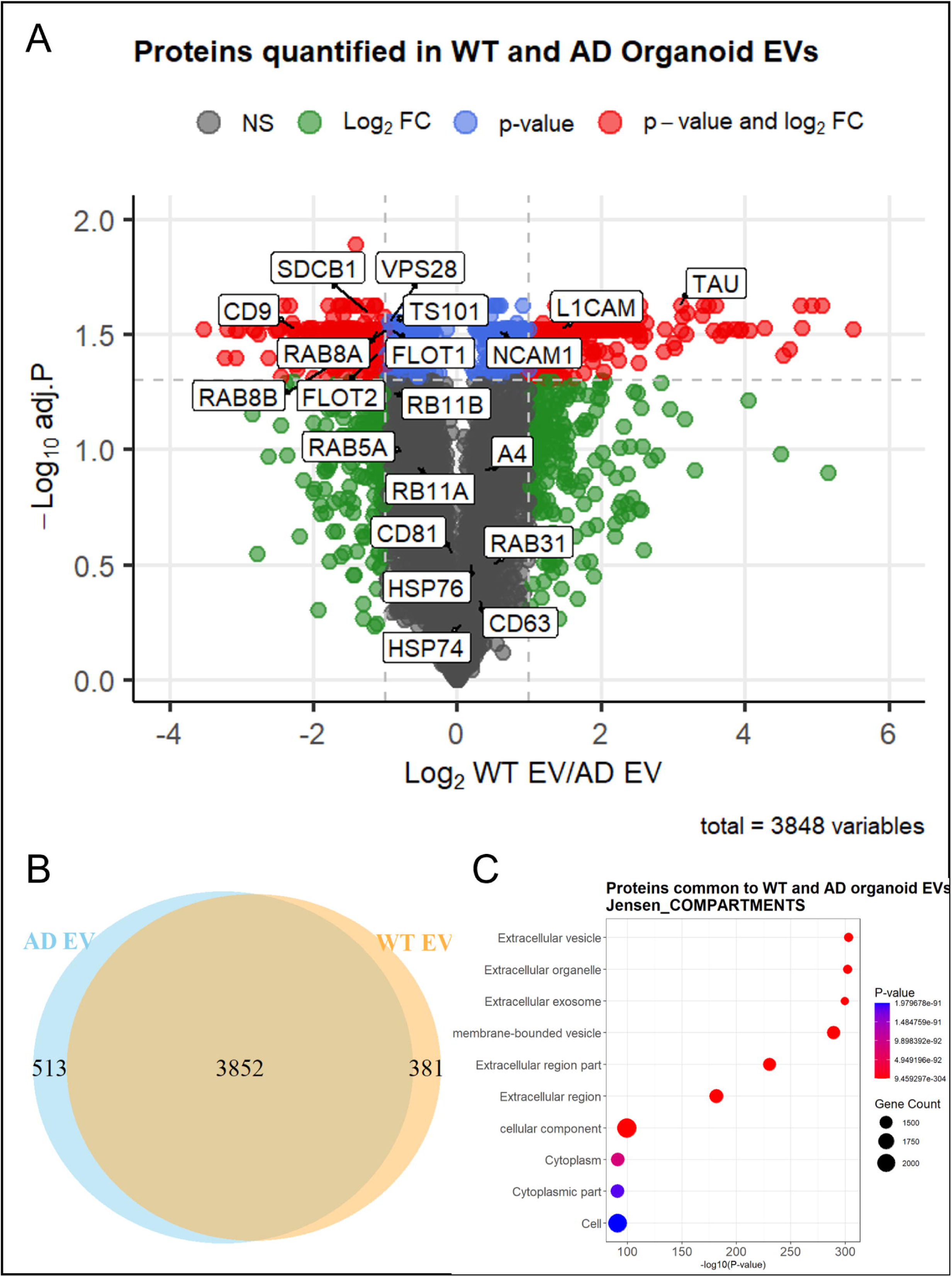
Deep proteomic profiling of organoid-derived EVs. Deep proteomic profiling of organoid-derived EVs. **A)** Volcano plot of proteins quantified in WT and AD organoid EVs and relative log2 ratio abundance comparison. EV protein markers as indicated: tetraspanins CD9, CD81, CD63; intraluminal proteins TS101 (TSG101), SDCB1, RAB5A, RAB31, VPS28; membrane lipid-raft proteins FLOT1/2; molecular chaperones (intraluminal) HSP74/76; brain-EV specific adhesion molecules NCAM1 and L1CAM; autophagy-related marker RAB8A, RAB8B, RB11A (RAB11A); Alzheimer’s disease-related proteins A4 (amyloid precursor protein; APP); TAU. Grey: non-significant; Green: FC only; Blue: adj p value only; Red: adj p and FC (significant; FC ≥ 2; adj p ≤ 0.05). **B)** Venn diagram represents proteins identified in both groups (3852), unique to AD EVs (513), and proteins unique to WT EVs (381). **C)** Enrichment analysis showing the top 10 enriched terms in the Jensen COMPARTMENTS database corresponding to proteins identified in both WT and AD organoid EVs. The top enriched terms include ‘Extracellular vesicle’, ‘extracellular exosome’, and ‘membrane-bounded vesicle’. Dot size corresponds to gene number; p-value is indicated on the x-axis and by color legend.

We further explored alternative secretory pathways in our proteomics data to determine whether the vesicle secretion dynamics observed in AD organoids could be attributed to a specific route of biogenesis, which could guide future studies targeting vesicle subpopulations. Comparative proteomics analysis of EVs revealed differences in autophagy and membrane trafficking machinery between WT and AD organoids. The amphisome marker MAP1LC3A (LC3A) was exclusively detected in WT EVs (3/3 replicates) and not detected in AD EVs (0/3 replicates), while MAP1LC3B showed 4.9-fold enrichment in WT EVs. Secretory autophagy regulators RAB8A and RAB8B, which mediate amphisome-plasma membrane fusion (Zubkova *et al*. 2024) were 2.2-fold and 2.0-fold more abundant in AD EVs (both adj. *p* = 0.030). The secretory SNARE protein STX3 was detected only in AD EVs (Kumar *et al*. 2018), while the degradative autophagy SNARE STX7 showed 1.5-fold enrichment in WT EVs (adj. *p* = 0.036) (Matsui *et al*. 2018). The lysosomal marker LAMP1 was 1.5-fold enriched in WT EVs (adj. *p* = 0.044). The classical EV marker CD9 and the ESCRT-I components VPS28, TSG101, and SDCB1 showed significant enrichment in AD EVs (4.8-fold, 1.9-fold, 1.8-fold, and 2.3-fold, respectively; adj. *p* = 0.030, 0.027, 0.028, and 0.026, respectively), indicating enhancement of conventional exosome biogenesis despite impaired amphisome secretion. Core autophagy proteins ATG5 and ATG7, while below the fold-change threshold, showed modest enrichment in AD EVs (1.3-fold, adj. *p* = 0.037 and 1.1-fold, respectively).

For qualitative analysis, proteins were considered identified if detected in ≥ 2 out of 3 replicates, resulting in 3852 proteins identified in both AD and WT EVs, with 513 proteins classified as unique to AD EV, and 381 unique to WT EV (**Figure 5B**). Enrichment analysis against the Jensen COMPARTMENTS database also showed significant enrichment of EV-related terms, including “Extracellular vesicle”, “extracellular exosome”, and “membrane-bounded vesicle” (**Figure 5C**). Visualization with principal component analysis (**Supplementary Figure 4**) using log2 intensities revealed dispersed clusters of replicates, with highest explained variance along the PC1 axis and separation of groups along the PC2 axis indicating close agreement between replicates within each condition.

### 3.5 Gene ontology: Enrichment analysis

Enrichment analysis with EnrichR identified several key pathways that were changed in proteins upregulated in either WT or AD EV samples (**Supplementary Figures 5-7**). AD organoid EVs were predominantly enriched in cell junction remodeling processes, such as cadherin binding (GO:0045296), cell-cell junctions (GO:0005911), focal adhesions (GO:0005925), and protein localization to plasma membrane (GO:0072659) (**Supplementary Figures 5-7**). Amyloid-beta binding activity (GO:0001540) was specifically enriched in AD organoid EVs, suggesting that EVs may play an active role in binding to pathological protein aggregates or are involved in their clearance (**Supplementary Figure 7**). Enrichment in cell migration regulation (GO:0030334) and cytoskeletal organization processes also indicate cellular remodeling in AD organoids (**Supplementary Figure 5**).

In contrast, WT organoid EVs showed strong enrichment in neuronal development processes, including axon guidance (GO:0007411), axonogenesis (GO:0007409), and nervous system development (GO:0007399) (**Supplementary Figures 5-7**). The most significant cellular compartment enrichments were collagen-containing extracellular matrix (GO:0062023) and neuron projection (GO:0043005) (**Supplementary Figure 6**), and enriched molecular functions included Protease Binding (GO:0002020), tubulin binding (GO:0015631), and calcium ion binding (GO:0005509), and neurotransmitter/receptor activities including Syntaxin-1 binding (GO:0017075) and acetylcholine receptor binding (GO:0033130) (**Supplementary Figure 7**).

These findings indicate that AD organoid EVs contain an increased abundance of proteins involved in pathological cell adhesion remodeling and amyloid interactions, while proteins upregulated in WT organoid EVs contain machinery supporting normal neuronal development and ECM organization, suggesting fundamental differences in the cellular processes being transmitted through vesicle-mediated intercellular communication.

### 3.6 Amyloid detecting dyes

Though there have been several studies showing that EVs can enhance the aggregation propensity of amyloidogenic proteins (Mirdha 2024; Budvytyte & Valincius 2023), the binding properties of EVs to amyloid-specific dyes have not been well characterized. The most well-known amyloid binding dye is ThT and ThT binding assays are the gold standard assay for monitoring amyloid formation in vitro (Biancalana & Koide 2010). EV purification samples were collected and added to the wells of a microplate with ThT. There was stable and high ThT fluorescence in the EV isolates from both WT and AD organoids throughout the time course of the assay (**Figure 6A**). The signal in the EV isolates was approximately 25 times higher than a control condition that contained synthetic liposomes of the same approximate size and that are made to have a lipid composition similar to the mammalian cell membrane (**Figure 6A**). ThT is known to bind to amyloid fibrils or cross-β-sheet aggregates that have either a fibrous or amorphous morphology but can also detect other biological polymers like RNA (Sugimoto *et al*. 2015). To provide further evidence for the presence of amyloids in EV isolates, we collected EVs and stained them with Amytracker 680, a conjugated oligothiophene dye known to selectively bind to amyloid fibril structures (Klingstedt *et al*. 2013). We visualized the stained EVs using STED microscopy and produced super-resolution images of EVs stained with AmyTracker 680 of both WT (**Figure 6B**) and AD (**Figure 6C**) EVs. Interestingly, AmyTracker 680 was quite responsive to the 775nm depletion laser used for STED imaging, allowing sub-diffraction limited imaging. The fields of view contained many EV-sized puncta of varying fluorescent intensity, suggesting some EVs contained more aggregated proteins than others (**Figure 6B and C**). A line profile intensity FWHK measurement of a representative AmyTracker 680 puncta was consistent with the size of EVs measured in the vFC and Cryo-TEM data (**Figure 2C**, **Figure 4H, and Figure 6B and C**). Amyloid proteins like Aβ_42_ and tau that were detected in EV isolates by WB and ELISA (**Figure 2**) could be the source of ThT and AmyTracker fluorescence in the EVs.

**Figure 6.**
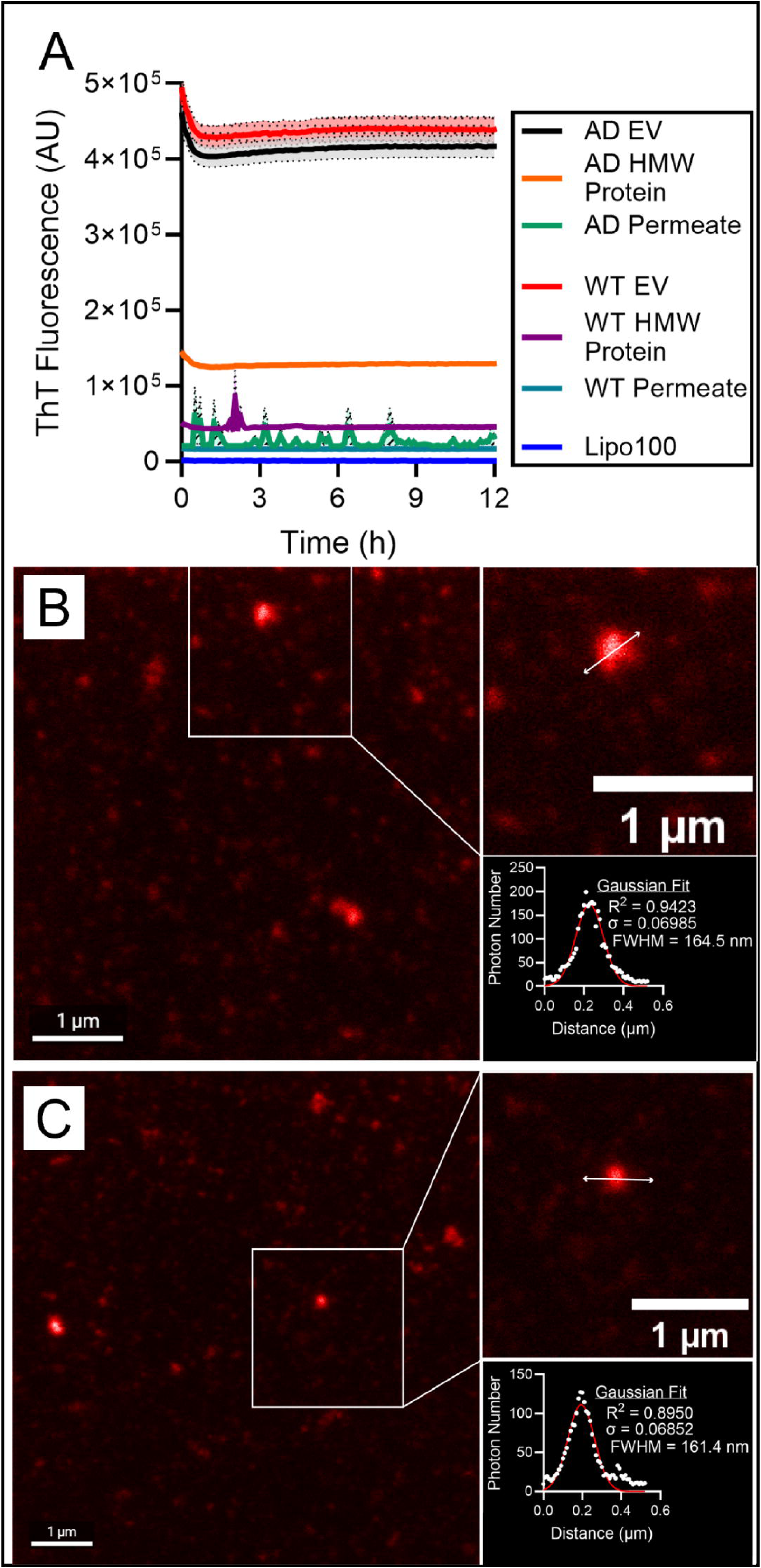
AD and WT organoid-derived EVs bind to amyloid detecting dyes. **A)** A ThT binding assay was performed to monitor amyloid formation in EV purification samples including an EV-depleted permeate control, a high MW Protein sample from the SEC column, and the final EV containing isolate. A synthetic liposome (Lipo100) control was also included. EV isolates purified from **B)** WT and **C)** AD conditioned media were stained with Amytracker 680 and imaged using STED / Confocal microscopy. The fluorescence gathered from a representative punctum from each condition was fit with a Gaussian curve and the vesicle diameter was estimated based on a FWHM measurement.

### 3.7 EVs can transfer from AD to WT organoids

EVs are a common method of cell-to-cell communication and are naturally secreted and endocytosed. The EVs purified from AD and WT organoid conditioned media contain amyloidogenic proteins and bind to amyloid specific dyes (**Figures 1 and 6**). Therefore, we theorized that EVs could pass from AD to WT organoids and bring with them their amyloid / amyloidogenic cargo. We stained AD organoids with Aco-650, a fluorogenic dye that is incorporated into endosomes of treated cells (**Figure 7A**). Aco-650 stained AD organoids were then rinsed several times and co-cultured with WT organoids over a 7-day period, where the two organoids were maintained in the same well of a 6-well culture plate with several full media changes. Notably, on the seventh day, the WT and AD organoids were imaged, revealing that Aco-650 signal was present in the WT organoids, suggesting that some endosomal material transferred from the AD organoid to the WT organoid (**Figure 7B**). All organoids were then fixed and underwent tissue clearing, a delipidation method that removed occluding lipids allowing for high resolution microscopy in thick 3D tissues (Hama *et al*. 2015). We immunostained all organoids with Aβ XP and Hoechst nuclear stain. Abeta XP puncta were present in all WT and AD organoids that were imaged, though there was substantially more signal present in the AD organoid, which is in accordance with previous observations of Aβ production in cerebrocortical organoids with the PSEN1 mutation (Labra *et al*. 2026) (**Figure 7C**). Strikingly, in the WT+AD co-culture, the WT organoids showed a significantly increased number of intratissue Aβ XP puncta as compared to the WT+WT cocultured organoids (**Figure 7C and D**). These data together suggest that co-culturing a WT organoid with an isogenic AD mutant organoid can result in the transfer of endosome material and potentially pathogenic amyloid-β peptides from one tissue to the other.

**Figure 7.**
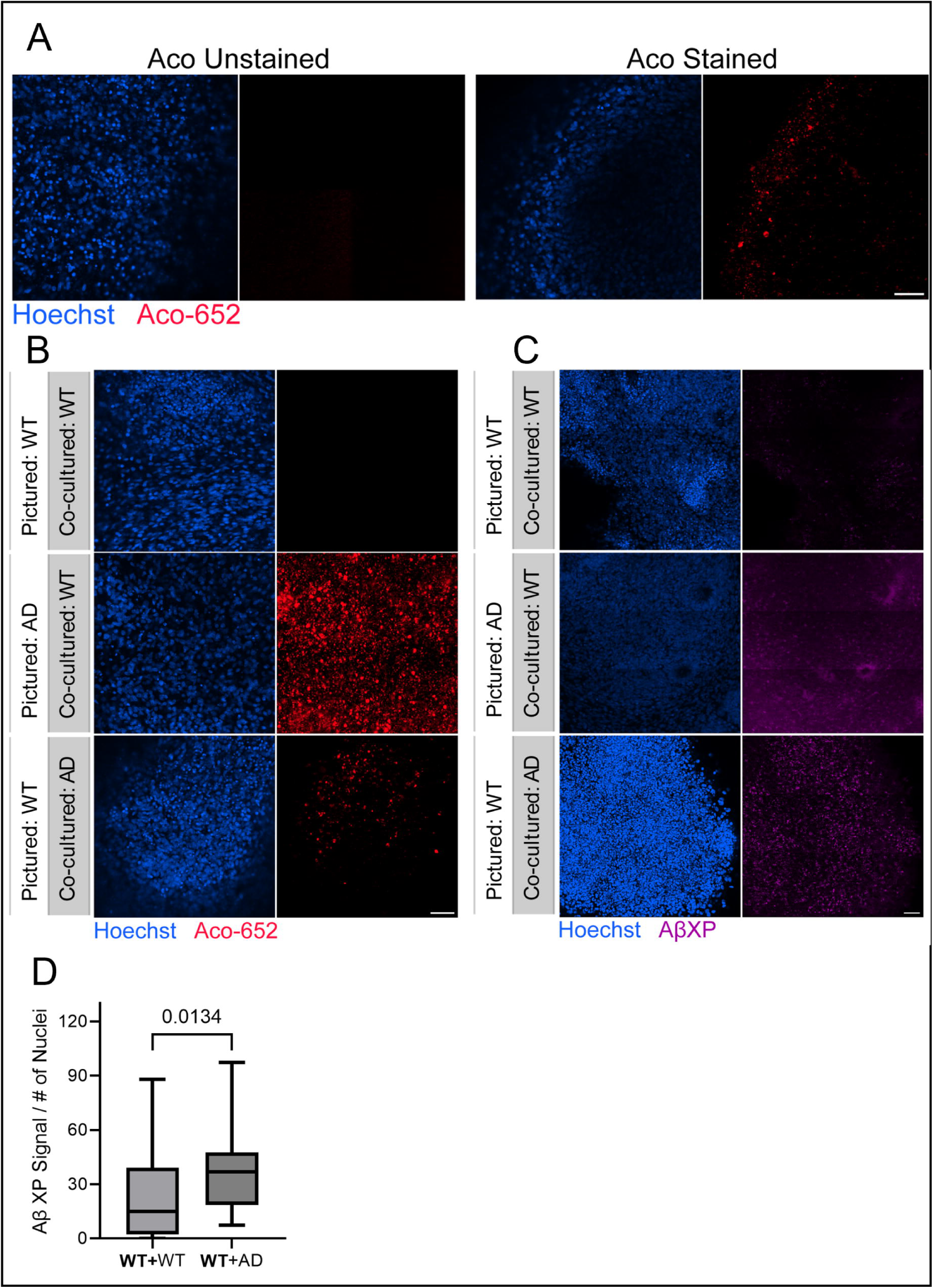
Intratissue Amyloid-β signal increases in WT organoids after one week co-culture with AD organoids. **A)** Live-cell imaging of an AD organoid stained with endosome marker and fluorogenic probe Aco-650 that intercalates into endosome membranes. WT organoids were co-cultured for one week with either another WT organoid or an Aco-650 stained AD organoid. **B)** On the seventh day, we performed live-cell imaging on representative organoids from each co-culture. **C)** After the live-cell imaging, all organoids were cleared and immunostained with primary antibodies that detect Aβ peptides. All scale bars represent 50 µm. **D)** Quantification showing increase in cumulative Aβ XP segment volume per cell in WT organoids after co-culture with an AD organoid (**WT+**AD) compared to a WT organoid after coculture with another WT organoid (**WT+**WT). Box plots show the median (line), the interquartile range (box), and whiskers extending to min/max data points. n = 27 images for **WT+**AD and n= 49 images for **WT+**WT. Statistical significance was assessed via student’s t test; p-value indicated above.

## 4. DISCUSSION

### 4.1 Enriching EVs from organoid conditioned media

EVs are valuable biological analytes that can be readily collected from organoid conditioned media. Publications within the last two years have described EVs secreted by cerebrocortical organoids, primarily focusing on transcriptomic and proteomic analyses (Liu *et al*. 2025a; Liu *et al*. 2025b; Silver *et al*. 2025; Forero *et al*. 2024). Because EVs reflect the health of their cells of origin, their contents can correlate with disease (Colombo *et al*. 2014). Cerebrocortical organoids offer several advantages as an EV source for studying AD compared with previously used sources like blood plasma, CSF, or postmortem brain homogenates (Wang *et al*. 2017; Fiandaca *et al*. 2015; Sardar Sinha *et al*. 2018). In our system, organoids were differentiated from an isogenic set of hiPSCs allowing us to isolate the effects of a single amino-acid substitution (PSEN1^M146V^) on AD-related phenotypes without genetic variability. Organoid culture media provides a continual supply of vesicles, unlike patient-derived sources that require venipuncture, lumbar puncture, or tissue processing. Lastly, since organoids remain viable for months (Wang 2018; Bubnys & Tsai 2022), they enable longitudinal EV collection as neurodegenerative features develop, making them a practical and biologically relevant model for examining AD-associated changes over time.

### 4.2 EV characteristics shared across genotype

Aβ_42_ and tau were both present in EV isolates immunoprecipitated with capture antibodies recognizing common EV membrane TS proteins (**Figure 2B**). ELISAs showed that Aβ_42_ was enriched by the TS protein IP and released into the eluate after EV lysis, indicating that Aβ_42_ is contained within TS EVs (**Figure 2C**). Aβ_42_ was also detected in the TS permeate, suggesting that this TFF fraction contains trace EVs that become enriched upon IP capture (**Supplementary Figure 2A**). Interestingly, tau did not copurify with Aβ_42_ (**Figure 2D**). Although tau was enriched in both WT and AD EV isolates relative to EV-depleted permeates (**Supplementary Figure 2B**), and notably Tau was found almost exclusively in the IP flow-through rather than the eluates (**Figure 2D**). Tau may appear in the flow-through either because it weakly associates with TS EVs and is not efficiently captured or alternatively is carried within EVs lacking TS proteins. As TS expression varies among EV subpopulations (Han *et al*. 2021), using other capture antibodies such as CD63 or LC3B could further resolve distinct EV subsets carrying tau (Visnovitz *et al*. 2025).

Vesicle flow cytometry (vFC) measures EV number, size, and phenotype (via immunofluorescence) at the single vesicle level (Sandau *et al*. 2020; Tekkatte *et al*. 2023). Used here, vFC revealed that fractionated AD organoid media contained about twice as many EVs as WT media. Regardless of the source, the EVs had similar sizes (range ∼85-270 nm, median 124 nm) and phenotype (TS expression at or below the LOD of ∼38 antibody molecules per vesicle; **Figure 3**). We did not measure TS expression on the cells, but EVs originate from different sub-cellular locations, and while EVs that originate from the cell surface and endosomal systems often carry TS, many EVs do not (Tekkatte *et al*. 2023; Oh *et al*. 2022), possibly because they originate from other subcellular locations, including autophagosomes and secretory lysosomes where TS is not abundant.

Thioflavin T and AmyTracker are amyloid specific dyes that bind to WT and AD EVs, and their signal is observable both in bulk fluorescence assays and by super-resolution microscopy. The interaction between Aβ and cellular membranes has been widely examined, with many studies reporting membranes enabling amyloid aggregation (Mirdha 2024; Budvytyte & Valincius 2023). Moreover, multiple papers have detailed how EVs influence the aggregation propensity of amyloidogenic proteins (Yuyama *et al*. 2012; Sanguanini *et al*. 2020; Halipi *et al*. 2024). Herein we show that both WT and AD organoid-derived EVs produced a markedly elevated ThT signal (about 25-fold over buffer control) in a standard ThT assay (**Figure 6A**), suggesting that components of the EV isolate, rather than the HMW protein fraction, contribute to this signal. Both the WT and AD organoid-derived EVs also fluoresce upon treatment with a different amyloid fibril binding dye, a conjugated oligothiophene called AmyTracker 680 (Klingstedt *et al*. 2013). STED super-resolution microscopy utilizes an excitation and depletion laser to achieve resolutions capable of resolving individual vesicles (Willig *et al*. 2006). AmyTracker 680 responded well to the depletion laser, and the resulting images showed vesicle□sized fluorescent puncta (**Figure 6B and C**). The vesicles varied in brightness, suggesting differences in amyloid aggregate load among EVs. A similar pattern was observed in our Cryo-TEM images, where vesicles displayed a range of luminal densities.

### 4.3 AD EV characteristics

There was surprising heterogeneity between AD and WT EV populations when visualized by Cryo-TEM. The AD EV isolates contained more MLVs that have several concentric membranes than the WT EVs (**Figure 4G and I**). MLVs have previously been observed in neurodegenerative contexts, including in EVs purified from PD patient CSF (Emelyanov *et al*. 2020). Notably, structures with fibril morphology were present within or around the larger AD EVs, whereas no comparable material was detected in WT isolates (**Figure 4G**). Additionally, fibril collections were occasionally observed independently (**Figure 4G**). While 2D class averaging did not yield identifiable features, several micrographs contain fibrils with amyloid-like twists, resembling PHF-type tau (**Figure 4G**). Fowler and coworkers were the first to show tau filaments present within EVs purified from AD brain homogenates (Fowler *et al*. 2025). Additional work is needed to characterize the molecular identity of the fibril morphology observed in our AD EV isolates. Immunogold labeling and cryo-electron tomography could identify their protein composition and relevance to disease pathology.

Alzheimer’s organoid derived EV proteomic reveals changes in the levels of proteins involved in pathological cell adhesion remodeling as well as an enhancement in amyloid interacting proteins (**Supplementary Figure 5-7**), while WT organoid EV proteomics reveal machinery supporting normal neuronal development and ECM organization (**Supplementary Figure 5-7**). This suggests a fundamental difference in the cellular processes conveyed through vesicle-mediated intercellular communication. Upregulation of proteins linked to axonogenesis and calcium dysregulation has been previously observed in MS-based proteomic studies of human AD brain tissue (Johnson *et al*. 2022; Askenazi *et al*. 2023), supporting the ability of our EVs to recapitulate *in vivo* disease-associated pathways. Importantly, our model enables characterization of extracellular signaling pathways in a complex AD brain organoid system that can be measured across time to mimic disease progression.

As AD is known to compromise autophagy–lysosome pathway function, we examined markers of secretory and degradative autophagy in WT and AD EVs, anticipating that autophagic extracellular vesicles (AEVs, <100 nm, (Mao *et al*. 2025)) would be present in our EV-enriched samples. Our proteomics data reveal a paradoxical autophagy phenotype in AD EVs: LC3A was undetected and LC3B was depleted 4.9-fold, indicating failed AEV formation, yet the molecular machinery required for secretory autophagy and AEV biogenesis was significantly enriched in AD EVs. Critically, RAB11B, the GTPase essential for MVB-autophagosome fusion and amphisome formation required for AEV biogenesis, was enriched 1.86-fold in AD EVs (adj.*p* = 0.056), indicating intact and upregulated amphisome formation. The presence of RAB11B, RAB8A/B, and STX3 – the complete molecular suite for secretory autophagy and AEV formation – without their conventional LC3-positive cargo reveals that the defect lies upstream of amphisome formation, most likely at the level of autophagosome biogenesis via ATG5/ATG7 (Lipinski *et al*. 2010; Itakura *et al*. 2012). This aligns with autophagy dysfunction reported in AD brain tissue, where ATG5 and ATG7 decline with aging and autophagosome–lysosome fusion is impaired (Nixon *et al*. 2005; Boland *et al*. 2008). AD organoids appear to mount a compensatory response by upregulating downstream secretory machinery, but without functional autophagosomes this machinery is aberrantly packaged into conventional CD9-enriched exosomes. This represents a novel extracellular biomarker signature of failed AEV biogenesis in AD: by the signature of RAB11, RAB8, and STX3 without LC3-positive cargo.

### 4.4 Transfer of membranes and A**β** between co-cultured organoids

Amyloid and tau pathologies are thought to spread throughout the central nervous system from initiating brain regions in AD progression (Brettschneider *et al*. 2015). Aβ is known to be present in EVs (Rajendran *et al*. 2006), and our results corroborate those observations (**Figure 2A and C**). Additionally, both WT and AD EVs bind to amyloid detecting dyes (**Figure 6**). Together, these findings suggest that EVs could serve as vehicles for the cell-to-cell transfer of Aβ aggregates.

A week-long co-culture of WT and AD organoids led to the transfer of endosome membranes between cultures and an increase in intra-tissue Aβ signal in the WT organoid. After one week of co-culture with an AD organoid, the WT organoids displayed Aco-650 puncta (**Figure 7B**) originating from the Aco-650–stained AD organoid. Therefore, the co-culture experiment shows direct transfer of endosome membranes from one culture to another. The same WT organoids had a significant increase in Aβ puncta after co-culture with an AD organoid compared to a co-culture with another WT organoid (**Figure 7C and D**). These correlated findings suggest that we may be observing the process of Aβ translocation between cells of the central nervous system and thus spread of amyloid pathology. Given the subsequent development of phosphorylated tau pathology in our PSEN1 mutant organoids (Labra *et al*. 2026), we attempted to perform the same signal transfer analysis with the AT8 antibody that detects phosphorylated tau, but we did not observe the same spread. It is possible that 8-to -10-week-old AD organoids are modeling the phase of AD that is characterized by the early spread of Aβ pathology, but not subsequent tau propagation.

## AUTHOR CONTRIBUTIONS

Conceptualization: AB; Data curation: NT; Formal Analysis: AB, NT, JS, JPN; Funding acquisition: NT, JRY, SAL, JWK; Investigation: AB, NT, JC, MA, AP, SC, KE, CB, CCK, JS, KV, KS; Methodology: AB, NT, SRL, SG, JS, JPN; Project administration: AB; Resources: JRY, JPN, SH, SAL, JWK; Software:; Supervision: AB, SRL, JRY, JPN, SH, SAL, JWK; Validation: AB, NT; Visualization: AB, NT, JS, JPN; Writing – original draft: AB, NT, JS, JPN; Writing – review & editing: AB, NT, SRL, JS, JRY, JPN, SAL, JWK

## ACKNOWLEDGMENTS

The authors would like to thank members of the Kelly and Lipton labs for their helpful comments and ideas for shaping this project. We’d like to particularly thank Emily P. Bentley who provided expert editorial support. Studies were funded by the NIH/NIA grants RF1AG061846, 5R01AG075862, 5R01AG077046 to JRY; T32 AA007456 to NT; R01 AG073418 to JWK; U01 AG088679, R01 AG056259, and R35 AG071734 to SAL.

## DATA AVAILABILLITY STATEMENT

The mass spectrometry proteomics data have been deposited to the ProteomeXchange Consortium via the PRIDE partner repository with the dataset identifier PXD076102. Analysis code is publicly available at: Turner, N.P. (2026). *Cerebrocortical EV analysis – WT vs AD* [RStudio]. GitHub. https://github.com/NataliePTurner/Cerebrocortical-EV-analysis---WT-vs-AD. All other data will be made available upon reasonable request to the corresponding authors.

