## Supplementary Figures for "Molecular and Structural Characterization Reveals Divergent Extracellular Vesicle Profiles Between Wild Type and Alzheimer’s Disease Cerebrocortical Organoids"

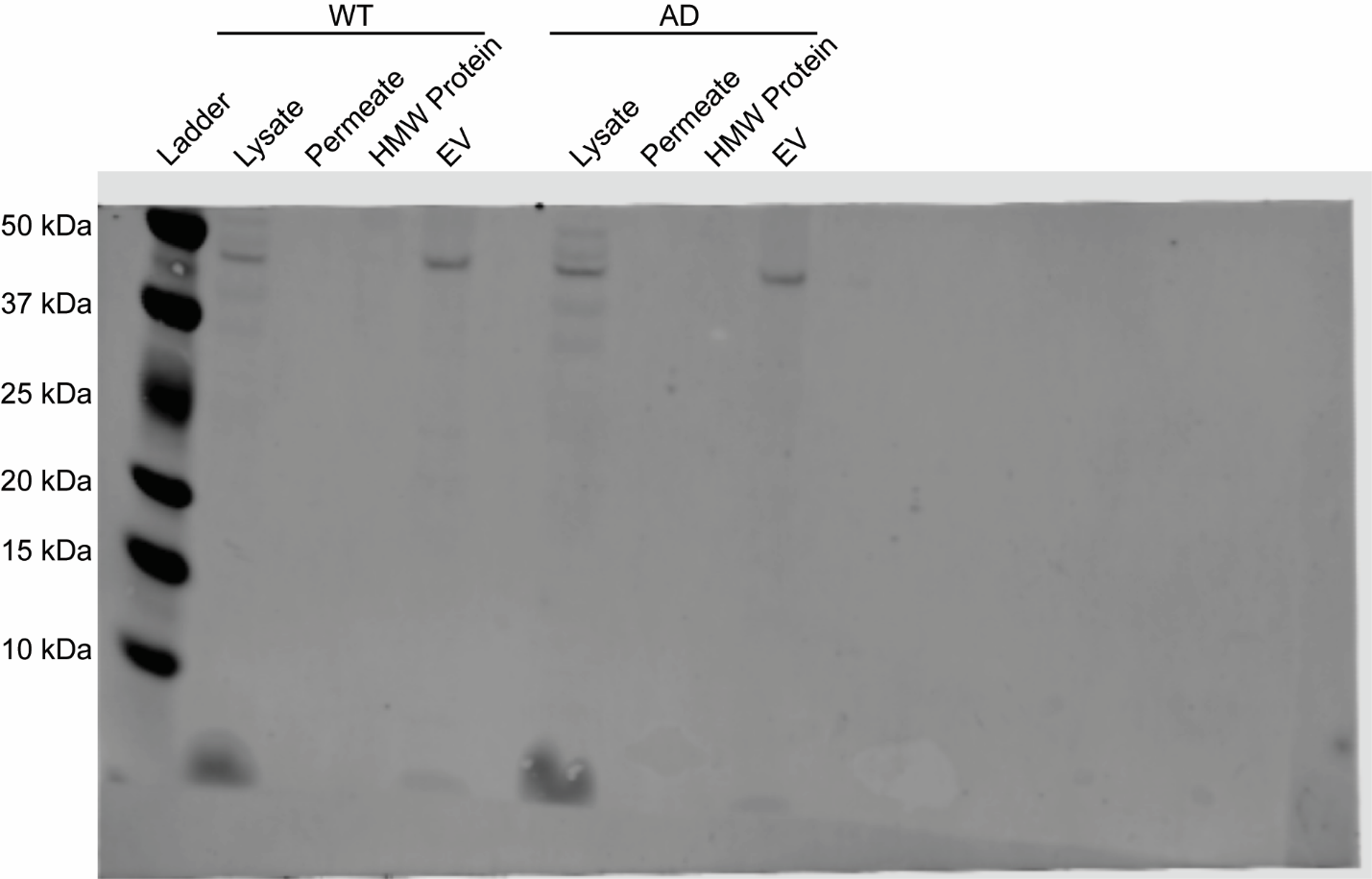

**Supplementary Figure 1. Full Aβ XP Western Blot.** Organoid conditioned media taken from 2-month-old WT and AD organoids was processes for EV enrichment, yielding an EV-depleted permeate, a HMW Protein sample, and a final EV isolate. Each purification sample and a sample of an organoid’s lysate was analyzed by Western Blot to detect amyloid-β peptides using the Aβ XP antibody.

A

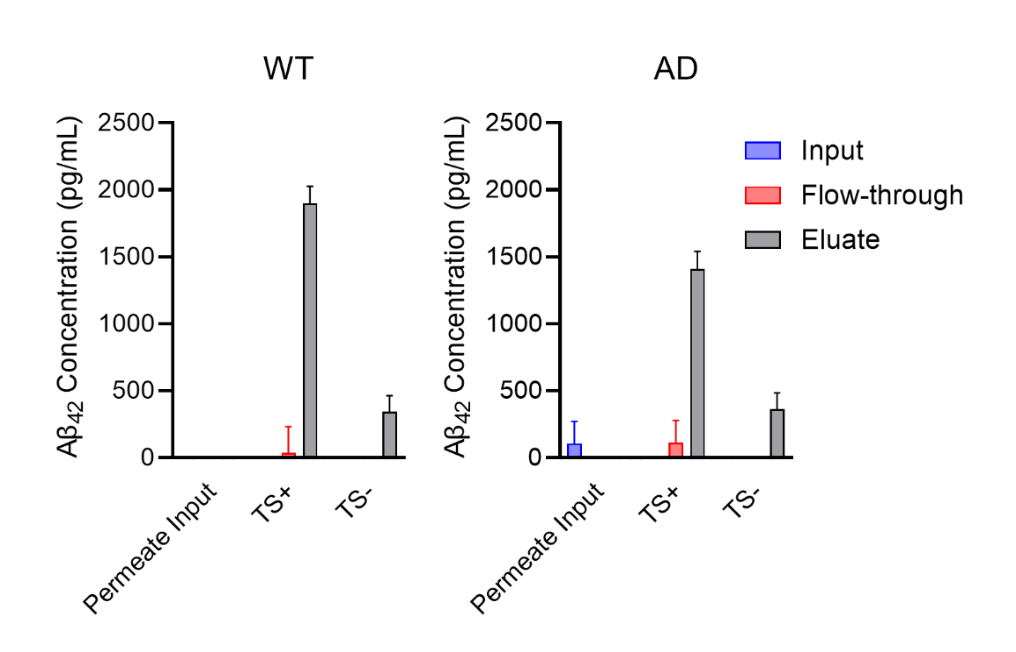

B

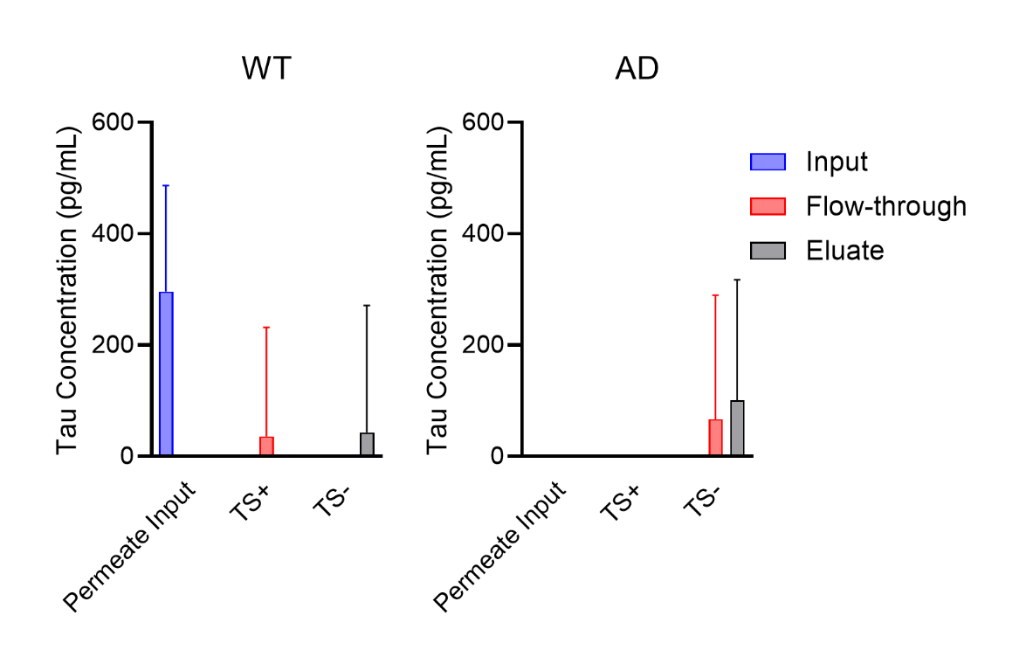

**Supplementary Figure 2. ELISA results detecting Aβ_42_ and tau in EV-depleted permeate samples.** Sandwich ELISAs were used to detect **A)** Aβ42 and **B)** tau (total) in IP samples taken from the EV-depleted permeates of both WT and AD organoid conditioned media. After IP, proteins that did not bind to the beads were labeled Flow-through and proteins associated with the beads were eluted from the beads using a detergent wash (1% SDS) and labeled eluate.

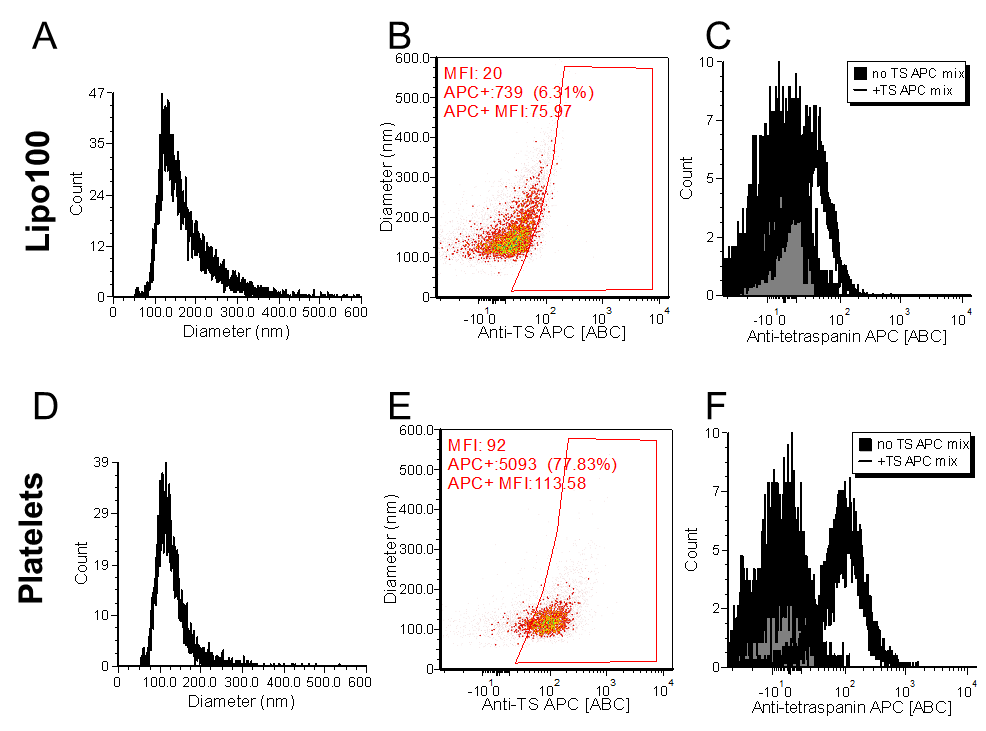

**Supplementary Figure 3. vFC control conditions. A and D)** Synthetic liposomes (Lipo100, top) and human platelet-derived EVs (bottom) were visualized by vFRed and APC fluorescence. **B and C)** Devoid of TS proteins, the Lipo100 liposomes exhibited extremely low levels of anti-TS antibody binding above background fluorescence, while **E and F)** platelet-derived EVs showed approximately 78% anti-TS antibody binding.

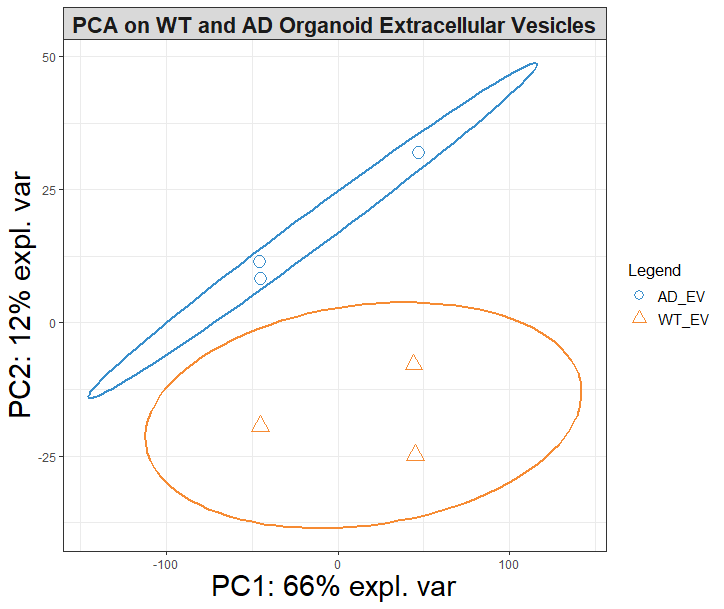

**Supplementary Figure 4. Principal component analysis (PCA) of log2‑transformed, median‑normalized protein abundances from WT (orange triangles) and AD EV samples (blue circles).** Each point represents a technical replicate. Circle-enclosed areas indicate 95% confidence ellipses for each experimental condition. Percent variance explained by PC1 and PC2 is indicated on the axes.

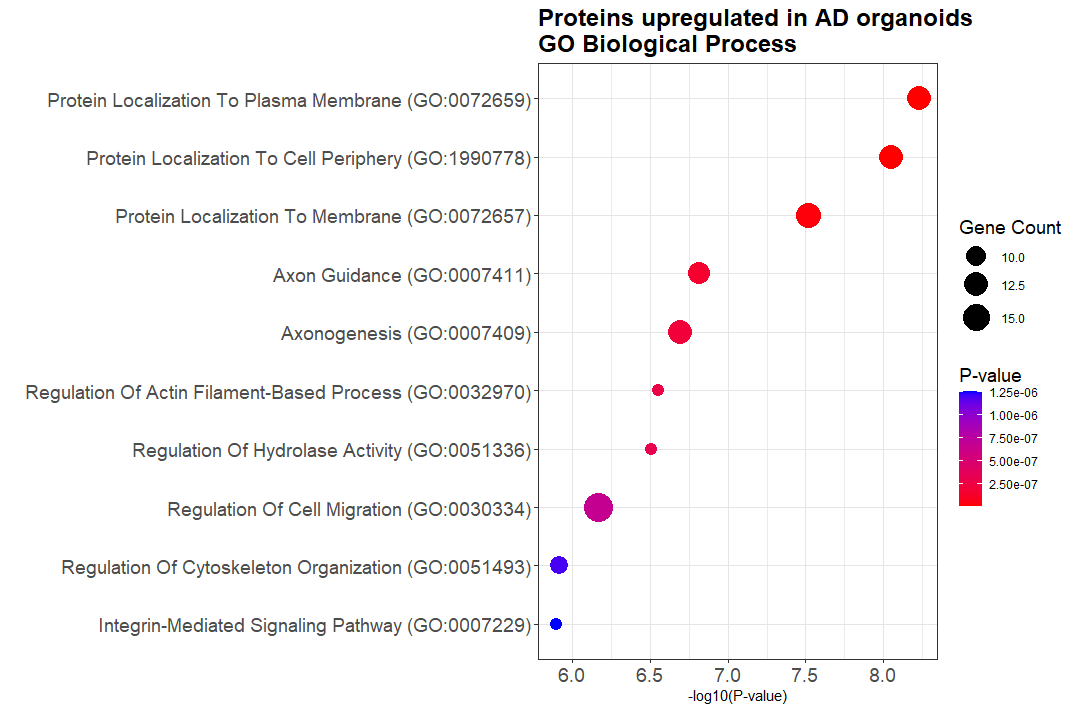

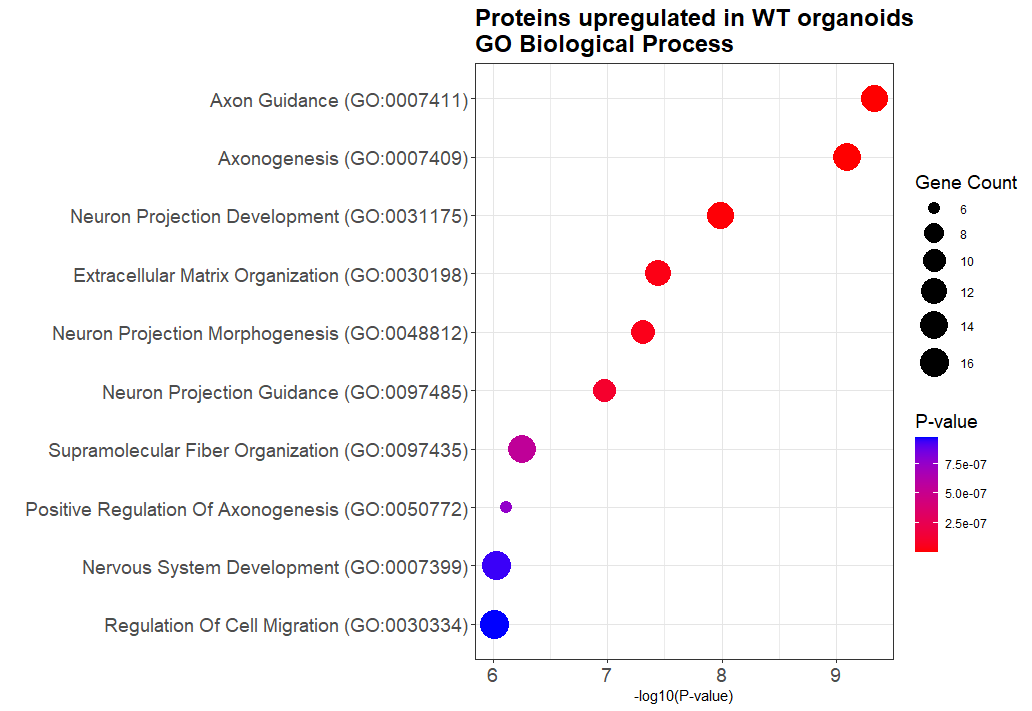
**Supplementary Figure 5. Gene Ontology enrichment analysis of biological process terms for proteins upregulated in WT or AD organoid-derived EVs**. Enrichment analysis was performed using Enrichr, and the top GO_Biological_Process categories are displayed for each condition. Bar width represents the gene count for each term, and color shading indicates the adjusted p-value. Terms are ranked by statistical significance, and ontology IDs are provided alongside descriptive labels.

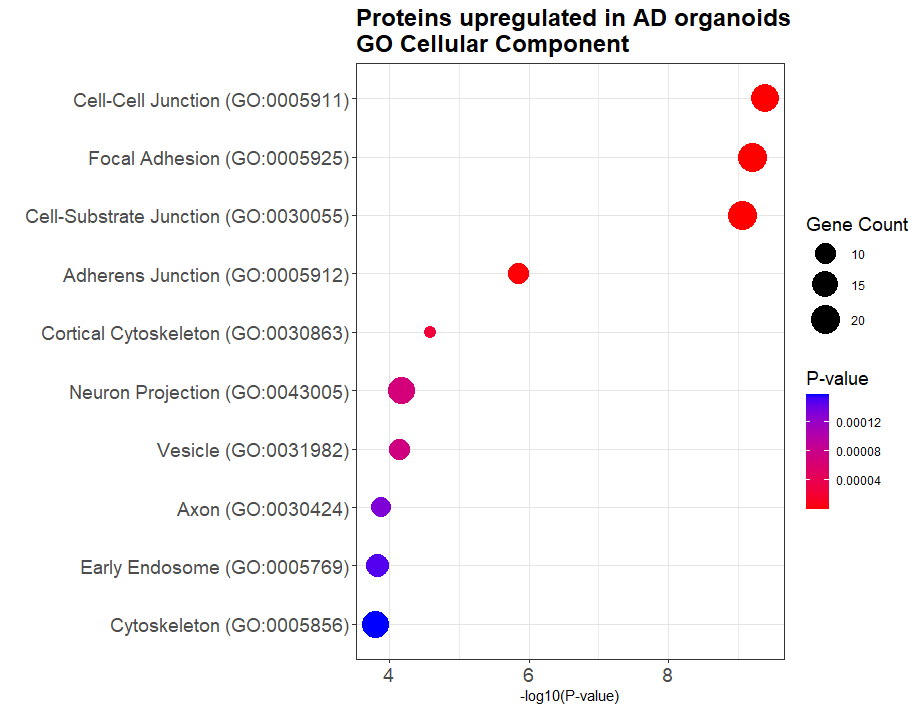

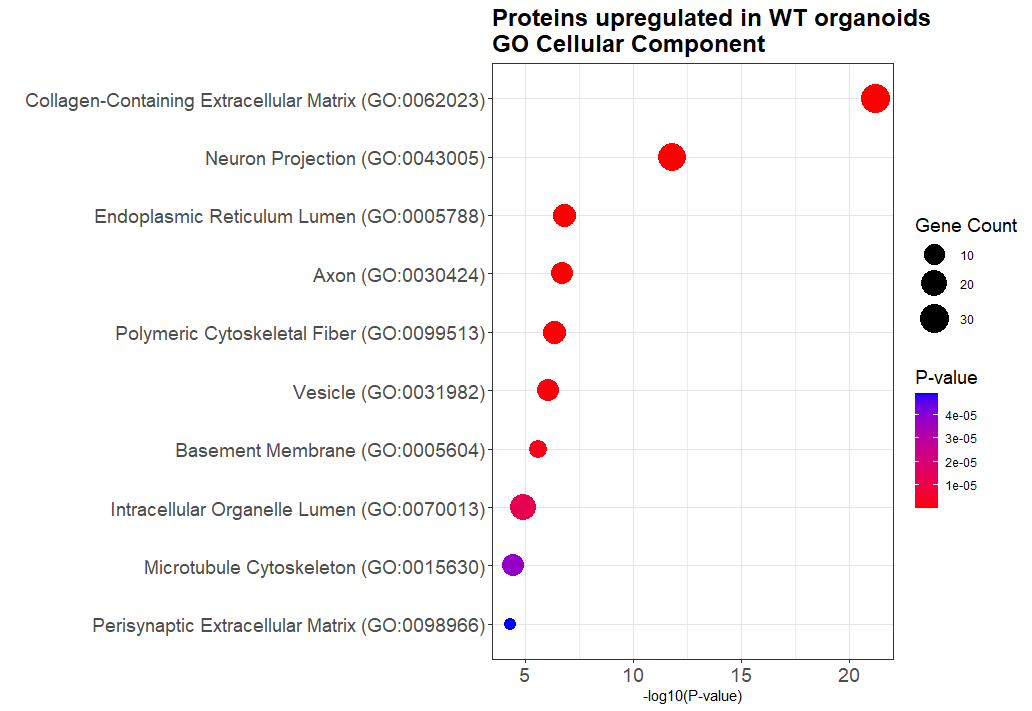
**Supplementary Figure 6. Gene Ontology enrichment analysis of cellular compartment terms for proteins upregulated in WT or AD organoid-derived EVs**. Enrichment analysis was performed using Enrichr, and the top GO_Cellular_Compartment categories are displayed for each condition. Bar width represents the gene count for each term, and color shading indicates the adjusted p-value. Terms are ranked by statistical significance, and ontology IDs are provided alongside descriptive labels.

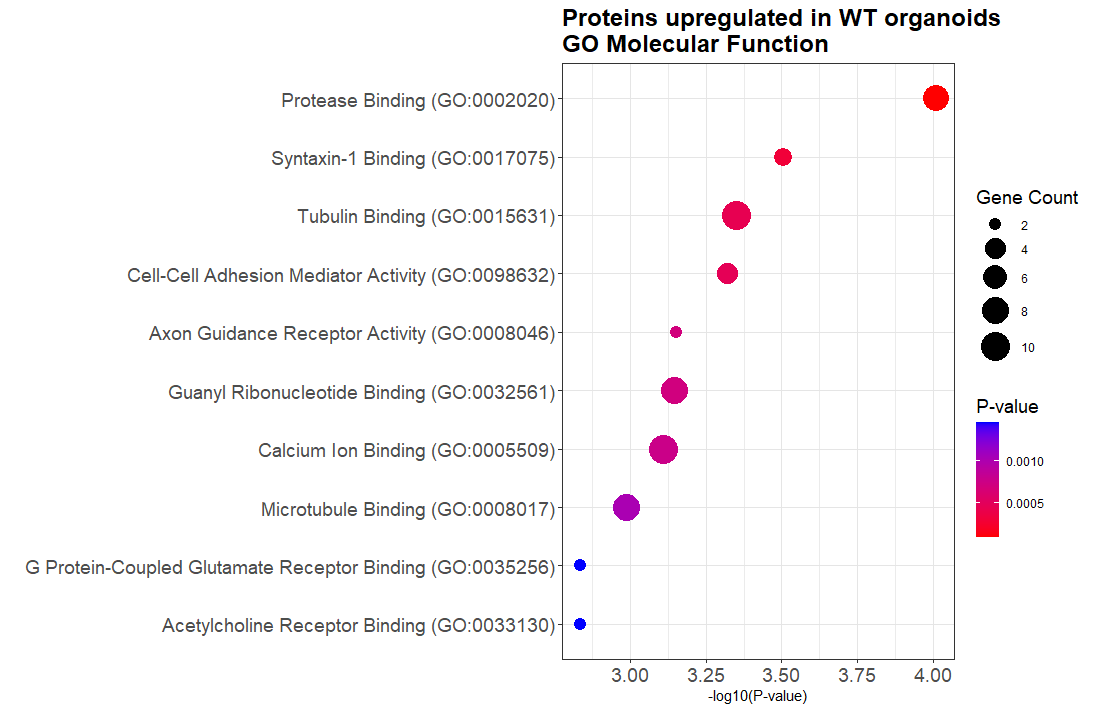

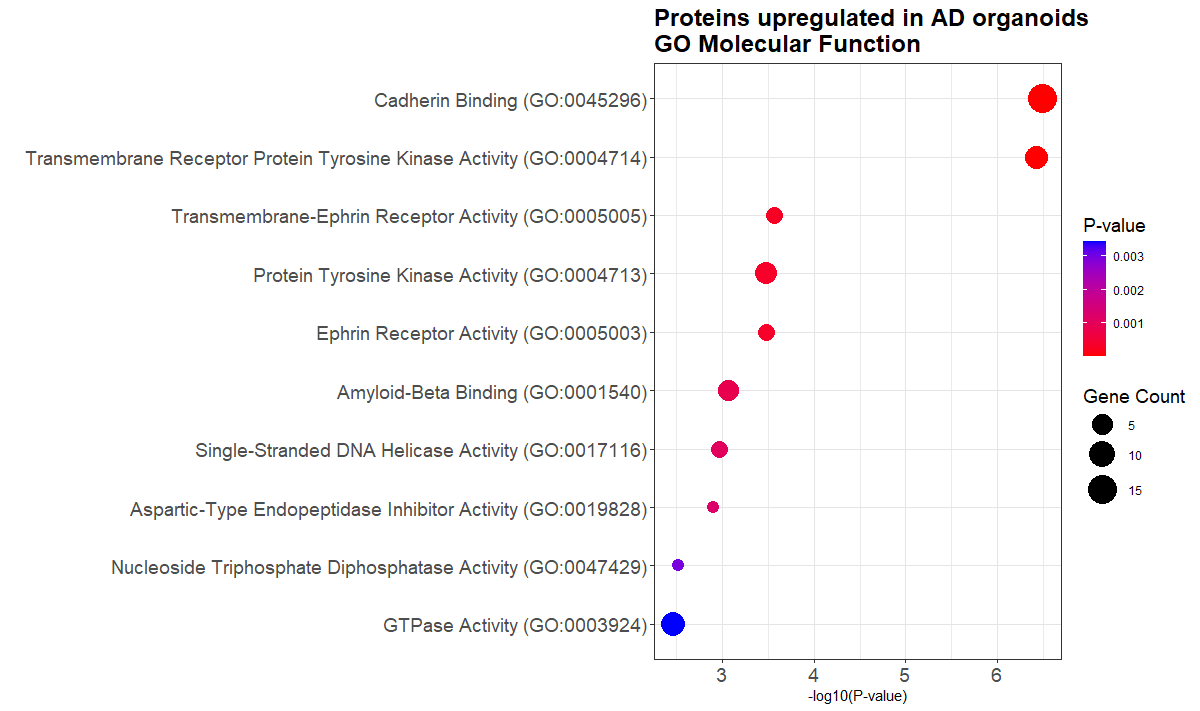
**Supplementary Figure 7. Gene Ontology enrichment analysis of molecular function terms for proteins upregulated in WT or AD organoid-derived EVs**. Enrichment analysis was performed using Enrichr, and the top GO_Molecular_Function categories are displayed for each condition. Bar width represents the gene count for each term, and color shading indicates the adjusted p-value. Terms are ranked by statistical significance, and ontology IDs are provided alongside descriptive labels.
